# The expansion and diversification of epigenetic regulatory networks underpins major transitions in the evolution of land plants

**DOI:** 10.1101/2024.09.22.614159

**Authors:** Romy Petroll, Ranjith K. Papareddy, Rafal Krela, Alice Laigle, Quentin Riviere, Kateřina Bišová, Iva Mozgová, Michael Borg

## Abstract

Epigenetic silencing is essential for regulating gene expression and cellular diversity in eukaryotes. While DNA and H3K9 methylation silence transposable elements (TEs), H3K27me3 marks deposited by the Polycomb repressive complex 2 (PRC2) silence varying proportions of TEs and genes across different lineages. Despite the major development role epigenetic silencing plays in multicellular eukaryotes, little is known about how epigenetic regulatory networks were shaped over evolutionary time. Here, we analyse epigenomes from diverse species across the green lineage to infer the chronological epigenetic recruitment of genes during land plant evolution. We first reveal the nature of plant heterochromatin in the unicellular chlorophyte microalga *Chlorella sorokiniana* and identify several genes marked with H3K27me3, highlighting the deep origin of PRC2-regulated genes in the green lineage. By incorporating genomic phylostratigraphy, we show how genes of differing evolutionary age occupy distinct epigenetic states in plants. While young genes tend to be silenced by H3K9 methylation, genes that emerged in land plants are preferentially marked with H3K27me3, some of which form part of a common network of PRC2-repressed genes across distantly-related species. Finally, we analyse the potential recruitment of PRC2 to plant H3K27me3 domains and identify conserved DNA-binding sites of ancient transcription factor (TF) families known to interact with PRC2. Our findings shed light on the conservation and potential origin of epigenetic regulatory networks in the green lineage, while also providing insight into the evolutionary dynamics and molecular triggers that underlie the adaptation and elaboration of epigenetic regulation, laying the groundwork for its future consideration in other eukaryotic lineages.

## Introduction

A key aspect of complex multicellularity involves the developmental regulation of gene expression in space and time. Spatiotemporal regulation is orchestrated by various regulatory genes, including transcription factors (TFs) and cell signalling pathways, but also involves regulation at the level of the epigenome (Zeitlinger and Stark 2010; Sparks et al. 2013). Epigenetic regulation reinforces cell identity in multicellular eukaryotes by dictating which parts of the genome should remain transcriptionally silent (Zhu and Reinberg 2011; Borg, Jiang, et al. 2021). As a consequence, epigenetic patterns are highly variable between different cell types and are reconfigured during multicellular development to specify cellular and tissue differentiation (Reik et al. 2001; Kawashima and Berger 2014). In most eukaryotes, this is shaped by three major types of heritable epigenetic marks - the methylation of DNA cytosine bases or by methylation of lysines 9 and 27 on the tail of histone H3 (i.e., H3K9me and H3K27me). H3K27me3 marks form facultative heterochromatin and are deposited by PRC2 to silence key development genes (Loubiere et al. 2019; Baile et al. 2021; Blackledge and Klose 2021). In contrast, DNA and H3K9 methylation often act together in a feedback loop to form constitutive heterochromatin, which interfaces with specific classes of short-interfering RNAs (siRNAs) to silence transposable elements (TEs) and repeat-rich sequences (Feng et al. 2010; Du et al. 2015).

The conserved and ubiquitous nature of epigenetic silencing across the eukaryotic tree of life has raised important questions about its evolutionary origins. While some studies have hypothesised that epigenetic silencing evolved to control invading “parasitic” elements like TEs and exogenous retroviruses (Déléris et al. 2021), others propose that TEs accumulate in eukaryotic genomes because of, not despite, epigenetic silencing mechanisms by suppressing homologous recombination of repetitive regions (Fedoroff 2012). PRC2 is likely to have emerged prior to the diversification of eukaryotes and consists of a conserved functional core of three-to-four subunits, E(z), ESC, Su(z)12 and p55 (Sharaf et al. 2022). In distant unicellular relatives of multicellular eukaryotes, PRC2 largely deposits H3K27me3 at TEs rather than genes, including in ciliates (*Paramecium;* Frapporti et al. 2019), in unicellular relatives of the red algae (*Cyanidioschyzon merolae;* (Mikulski et al. 2017; Hisanaga, Romani, et al. 2023) and in Stramenopiles (*Phaeodactylum tricornutum;* (Veluchamy et al. 2015; Zhao et al. 2021)). In the chlorophyte green microalga *Chlamydomonas reinhardtii*, a member of an early diverging lineage of the Viridiplantae phylum, H3K27me3 is not detectable despite presence of the H3K27 methyltransferase subunit E(z) (Shaver et al. 2010; Huang et al. 2017; Khan et al. 2018). Perturbed PRC2 activity in *Chlamydomonas* results in the derepression of some TEs (Shaver et al. 2010), suggesting that the ancestral role of PRC2 in TE silencing might also be conserved at the root of the Viridiplantae. However, a lack of genomic level data for H3K27 methylation in unicellular green algae has prevented definitive conclusions.

Because PRC2 plays such a key role in specifying cell differentiation in multicellular eukaryotes, its adaptation towards regulating gene expression could have been a major factor in facilitating the transition to multicellular life (Gombar et al. 2014). The emergence of multicellularity in plants, coupled with the transition to land and subsequent emergence of seed plants, were major events in plant evolution that all coincided with repeated bursts of *de novo* gene emergence, whole genome duplication and neo- functionalisation (de Vries and Archibald 2018; Bowman 2022; Barrera-Redondo et al. 2023). The emergent genetic novelty arising from these events has gained distinct spatial and temporal expression over evolutionary time. This is likely to have been facilitated by the adaptation and elaboration of epigenetic silencing mechanisms during the course of plant evolution, resulting in epigenetic landscapes where H3K9me2 and H3K27me3 largely repress either TEs or genes, respectively (Vigneau and Borg 2021). *Marchantia polymorpha* and *Anthoceros agrestis*, two extant representatives of the bryophyte lineage, still bear remnants of this progressive adaptation since PRC2 deposits H3K27me3 at both TEs and genes (Montgomery et al., 2020; Hisanaga et al., 2023). In flowering plants, H3K27me3 predominantly silences genes and is reprogrammed throughout the plant life cycle to facilitate development (Vigneau and Borg 2021), although its impact on TE silencing is also evident albeit to a lesser degree than at genes (Hisanaga, Romani, et al. 2023; Hure et al. 2025). H3K27me3-marked TEs lying in the vicinity of *Arabidopsis* genes contain several TF-binding sites, suggesting that their co-option may have facilitated the recruitment of PRC2 to genes during plant evolution (Hisanaga, Romani, et al. 2023). Studies in *Arabidopsis* also suggest that H3K9me2 silences a substantial number of pollen-specific genes, which are re-activated via epigenetic reprogramming in the vegetative cell (Borg, Papareddy, et al. 2021). Similar reprogramming of DNA and H3K9me2 also occurs in the central cell of the female gametophyte, suggesting that H3K9me2-mediated gene silencing may be a general feature of flowering plants (Pillot et al. 2010; Ibarra et al. 2012; Park et al. 2016). The extent to which H3K9me2 methylation silences genes in other plant species, however, remains unclear. This raises fundamental questions about how adaptations in epigenetic regulation could have contributed to the emergence of complex multicellularity during land plant evolution. At what time point in evolution did these epigenetic pathways recruit genes? What were the dynamics of this evolutionary adaptation and did this favour particular gene families? And are epigenetically-silenced genes common among all land plants or are they specific to each major clade? Such questions remain largely unresolved, not least through a systematic assessment across a major eukaryotic lineage.

Here, we perform a comparative analysis of epigenomes from six distantly related species of the Viridiplantae, including unicellular green algae, bryophytes and flowering plants. We provide insight into the nature of heterochromatin in the unicellular alga *Chlorella sorokiniana* (Trebouxiophyceae), a member of the chlorophyte lineage that diverged from the streptophyte lineage over 1 billion years ago (Leliaert et al. 2011; Moczydlowska et al. 2011). We show heterochromatin in *Chlorella sorokiniana* is defined by H3K27me3 but is partitioned into two distinct forms by the co-occurrence or absence of H3K9me2. A substantial number of *Chlorella sorokiniana* genes marked solely with H3K27me3 suggests that PRC2-regulated genes predate the emergence of land plants. By aging genes across the plant kingdom using genomic phylostratigraphy, we reveal how evolutionarily young genes are preferentially silenced with H3K9me2 in flowering plants, some of which become re-activated during reproductive development in *Arabidopsis*. In contrast, genes that emerged during and after Streptophyte evolution are preferentially silenced with H3K27me3, highlighting the preferential recruitment of PRC2 towards genes that emerged during these foundational periods of plant evolution. We also show how PRC2 silences a conserved gene network in the vegetative phase across land plants, which includes ancient regulators of the plant life cycle, architecture and reproductive development. Finally, we analyse *cis*-regulatory elements within H3K27me3 domains across distantly-related species and reveal prominent enrichment for known PRC2-interacting TFs, which may have helped establish some of the first PRC2-controlled gene networks in plants.

## Results

### H3K27me3 marks both TEs and genes in the chlorophyte green microalga *Chlorella sorokiniana*

The Viridiplantae (or green lineage) represents the most dominant phylum in the Archaeplastida kingdom and is represented by modern-day chlorophyte and streptophyte algae and their land plant relatives. Existing chromatin profiles from chlorophyte green algae are only available for the model species *Chlamydomonas reinhardtii* (Ngan et al. 2015; Strenkert et al. 2022), which lacks H3K27me3 and displays amino acid residue changes surrounding K27 on the tail of histone H3 (Shaver et al. 2010). To determine whether the alternative H3 sequence is common to other green microalgae, we performed a phylogenetic analysis of histone H3 variants from representatives of the Chlorophyta and Streptophyta, which diverged from each other *circa* 1250 mya (Evanovich et al. 2020). Interestingly, the characteristic S28T mutation and A29 deletion in *Chlamydomonas* H3 was only present in its close relative *Volvox carteri* (**Supplemental Fig. S1A**), suggesting that these residue changes are specific to the volvocine green algae within the Chlamydomonadales order. To clarify the presence and genomic distribution of H3K27me3 in chlorophytes, we selected *Chlorella sorokiniana* (from hereon in *Chlorella*) as a representative green unicellular alga, which belongs to the Trebouxiophyceae, a sister clade to the Chlorophyceae (**Fig. 1A**). Consistent with *Arabidopsis*-like H3.1- and H3.3-type tails (**Supplemental Fig. S1A**), we could detect both H3K4me3 and H3K27me3 by western blotting in *Chlorella* (**Supplemental Fig. S1B**).

**Figure 1.**
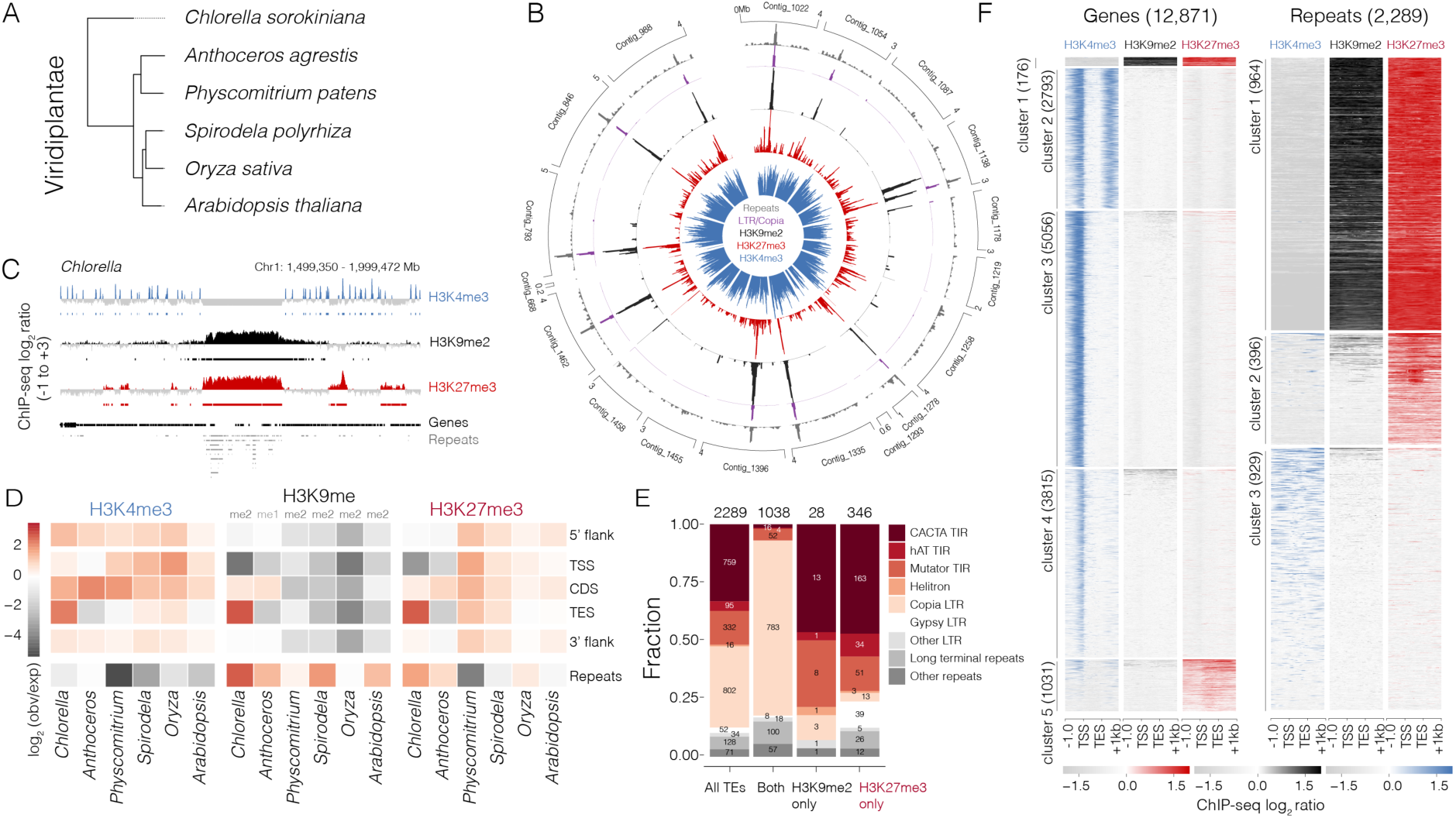
The heterochromatin landscape of the unicellular chlorophyte alga Chlorella sorokiniana. (A) Phylogenetic relationships of the six species analysed in this study. (B) Circos plot representing the density of repeats (grey), LTR/COPIA elements (purple) and ChIP-seq peaks of H3K9me2 (black), H3K27me3 (red) and H3K4me3 (blue) along each contig in the *Chlorella* genome. (C) ChIP-Seq tracks of the repeat-rich hotspot on chromosome 1. Coloured and grey shading indicate enriched or depleted signals, respectively. Coverage is represented as the ChIP-seq log_2_ ratio relative to H3. (D) Distribution of H3K4me3, H3K9me2 and H3K27me3 peaks at genomic features across the green lineage. The log_2_ enrichment or depletion is calculated relative to the frequency of genomic features in each species relative to 10,000 random permutations. TSS = translation start site, CDS = coding sequence, TES = translation end site. (E) Distribution of repeat classes marked with either H3K9me2, H3K27me3 or both alongside their relative frequency of all classes in the *Chlorella* genome. (F) Heatmaps of H3K4me3, H3K9me3 and H3K27me3 enrichment centred on genes and repeats in *Chlorella*. Plotted is the ChIP-seq log_2_ ratio relative to H3. ChIP-seq was performed with two biological replicates.

Using chromatin immunoprecipitation coupled with DNA sequencing (ChIP-seq), we obtained genomic profiles for the histone marks H3K4me3, H3K9me2 and H3K27me3 from *Chlorella* cells synchronised in G1 phase (**Fig. 1B-C; Supplemental Fig. S1C**). We observed that each histone mark was enriched at distinct genomic regions as in several land plants, suggesting similar features of chromatin in this unicellular green alga (**Fig. 1D**). Most *Chlorella* contigs were characterised by one or more repeat-rich hotspots strongly enriched for both H3K9me2 and H3K27me3 but depleted in active H3K4me3 marks (**Fig. 1B-C**). LTR/Copia elements were the only class of TEs present within these repeat-rich hotspots (**Fig. 1B**), suggesting that these regions represent the centromeres as reported in another strain of *Chlorella sorokiniana* (Wang et al., 2024). Consistently, H3K9me2 and H3K27me3 peaks were strongly enriched at repeats (permutation tests, log_2_ enrichment > 1; **Fig. 1D**) and were frequently deposited at the same repeats (cluster 1; 964 TEs), while H3K27me3 was enriched at a second group independently of H3K9me2 (cluster 2; 396 TEs) (**Fig. 1E-F**). Closer inspection revealed that the class of repeats present in these clusters also differed (**Fig. 1E**). Co-deposition of H3K9me2 and H3K27me3 mainly occurred at LTRs (77.9%; 809/1038), most prominently at LTR/Copia elements located in the putative centromeres (**Fig. 1B,E**) (Wang et al., 2024). In contrast, sole deposition of H3K27me3 mostly occurred at DNA transposons belonging to the CACTA and hAT terminal inverted repeat (TIR) element family (71.7%; 248/346), as well as Gypsy LTRs compared to the genome average (**Fig. 1E**). We also noted low levels of fragmented H3K4me3 deposition at a subset of repeats (cluster 3; 929 repeats) (**Fig. 1F**), which appears to be associated with their close proximity to the TSS of genes (**Supplemental Fig. 1D**). Of these, only 370 repeats (12% of total) overlapped with an H3K4me3 peak and were mostly (94%; 349/370) classified as TIR elements (**Supplemental Fig. 1E).** Whether these TE fragments represent exapted TE genes, which often originate from DNA transposons, remains to be determined (Hoen and Bureau 2015).

While H3K9me2 was largely restricted to the repeat-rich hotspots, H3K27me3 was dispersed across the chromosomes (**Fig. 1B**). Clustering of the chromatin profiles over genes revealed that H3K27me3 was also deposited at multiple genes in *Chlorella* (**Fig. 1F**, cluster 5: 1031 genes, 8% of gene models). Functional classification of these genes revealed an over-representation of 24 functional terms in the GO, KEGG and KOG classifications. They highlighted the targeting of transposon-encoded proteins and related processes, including DNA integration and RNA-dependent DNA replication, but were also related to protein phosphorylation, proteolysis and carbohydrate metabolism (**Supplemental Fig. S1F**). Thus, H3K27me3 does not appear to solely silence TEs in *Chlorella* but also marks protein-coding genes, suggesting that PRC2-mediated gene regulation likely predated the emergence of land plants.

## Evolutionarily young and old genes are silenced by distinct epigenetic states in plants

To gain insight into when epigenetic recruitment of genes occurred during land plant evolution, we re-analysed ChIP-seq datasets from five Viridiplantae species **(Fig. 1A; Supplemental Table S1)** - the two bryophytes *Physcomitrium patens* and *Anthoceros agrestis* and the three flowering plants *Spirodela polyrhiza*, *Oryza sativa* and *Arabidopsis thaliana* (from hereon in referred to by their respective genus). We focused on datasets derived from vegetative tissue of each species, namely thallus tissue for the bryophytes, mature leaf for *Oryza* and *Arabidopsis*, and whole minute plants for the duckweed *Spirodela*. Datasets from different studies were combined in the case of *Physcomitrium*, *Spirodela* and *Oryza* so as to include H3K9me2 profiles (**Supplemental Table S1**), which we also confirmed to be highly correlated across independent datasets **(Supplemental Figure 2A-B)**. We assigned phylostratigraphic ages to genes in each species then assessed the enrichment or depletion of genes marked with H3K9me1/2 and H3K27me3 in each phylogenetic rank (**Supplemental Fig. S2C; Supplemental Table S2**). Our analysis revealed a consistent and biased pattern that correlated with evolutionary age in divergent species of plants (**Fig. 2A-F**).

**Figure 2.**
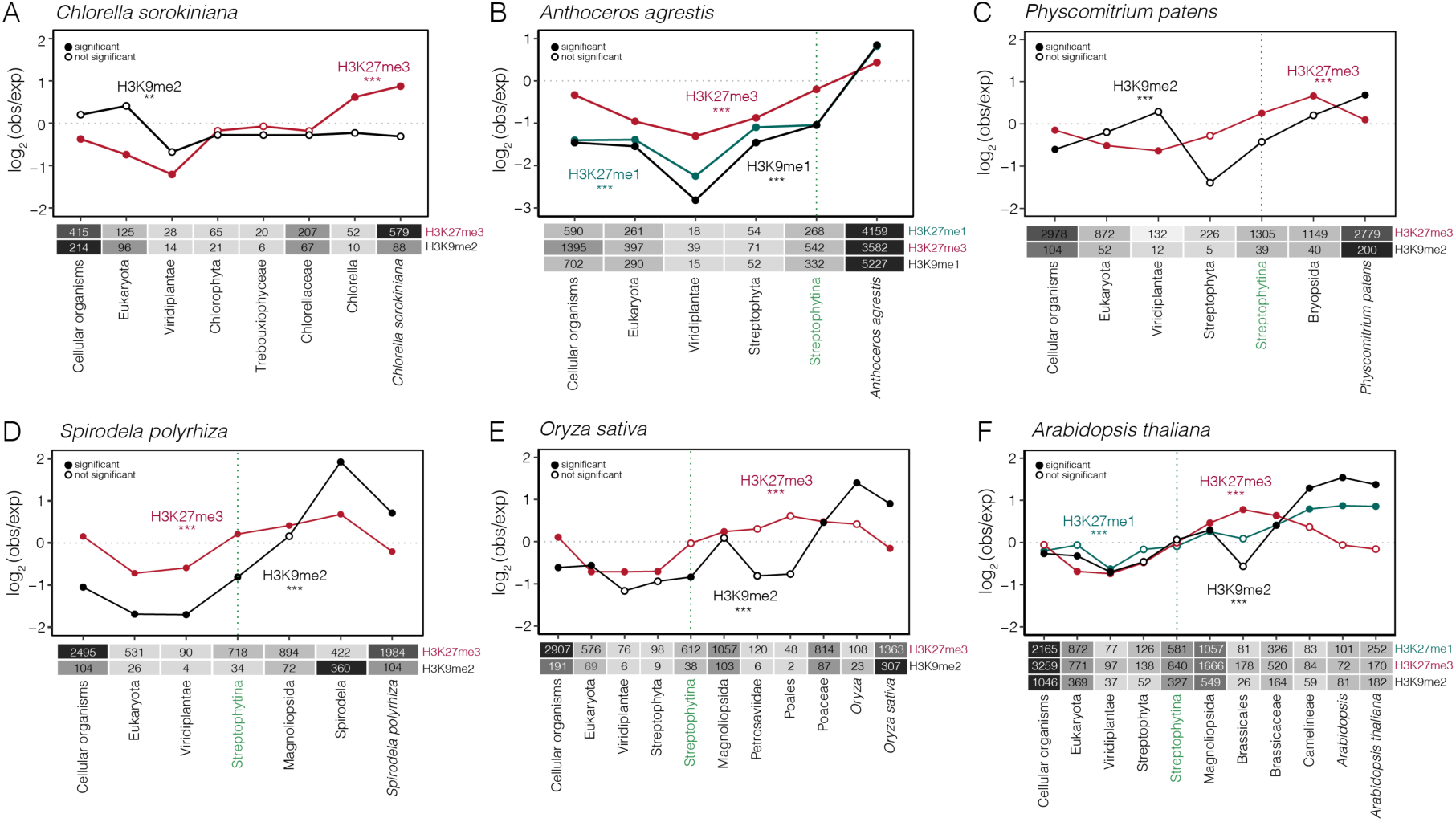
Epigenetic state correlates with the evolutionary age of genes across the Viridiplantae. Line plots showing the relative enrichment of H3K9me1/2 (black), H3K27me1 (green) and H3K27me3 (red) at genes grouped by their phylostratigraphic (or evolutionary) age in *Chlorella* (**A**), *Anthoceros* (**B**), *Physcomitrium* (**C**), *Spirodela* (**D**), *Oryza* (**E**) and *Arabidopsis* (**F**). Genes are ordered left-to-right from the oldest genes (i.e., those with homology in all cellular organisms) to the youngest (i.e., those restricted to each respective species). Plotted is the log_2_ ratio of marked genes relative to all genes in each phylogenetic rank. The green dotted line represents the Streptophytina rank when complex multicellularity arose during streptophyte evolution. The heatmap below each plot summarises the total number of genes marked with either epigenetic mark in each rank. A chi-square test was used to determine whether the distribution of phylogenetic ranks among the genes marked with each epigenetic mark was significantly different when compared to the genome average and is indicated under each mark (*** = *p* < 0.001). Solid dots indicate phylogenetic ranks that are significantly enriched or depleted as determined using a two-sided Fisher’s exact test with Bonferroni correction. Empty dots indicate no significance. See Supplementary Table 3 for a statistical summary relating to this figure.

Among the land plants, genes marked with H3K9me2 (or H3K9me1 in *Anthoceros*) were significantly enriched for taxonomically-restricted genes (TRGs) that have limited or untraceable homology in other species (**Fig. 2B-F; Supplemental Table S3**). This association was also evident at the level of DNA methylation, with TRGs having significantly higher rates of methylated cytosines in most contexts compared to older genes, particularly for CHH methylation (**Supplemental Fig. S3**). In *Anthoceros* and *Arabidopsis*, H3K27me1-marked genes were also significantly enriched for TRGs (**Fig. 2F**), further highlighting their tendency to be silenced with constitutive heterochromatin. To confirm whether the TRGs marked with H3K9me2 have any functional relevance in *Arabidopsis*, we assessed their expression pattern across development. We further calculated *tau* scores for each gene to determine whether these had a narrow (*tau* > 0.8) or broad (*tau* < 0.5) expression patterns (Lüleci and Yılmaz 2022). H3K9me2-marked older genes (ranks 1-6) had a narrow range of significantly higher *tau* scores compared with all *Arabidopsis* genes of the equivalent phylogenetic rank, suggesting restricted expression patterns during development (**Supplemental Fig. S4A-B**). TRGs specific to the Brassicaceae (ranks 7-10) also had a high and narrow range of *tau* scores and were less likely to be transcribed than older genes, which was more pronounced among those marked with H3K9me2 (**Supplemental Fig. S4A-B**). H3K9me2-marked TRGs tended to transcribe in reproductive stages like microspores, pollen and early zygotes (**Supplemental Fig. S4B**). No such bias was evident for H3K27me3-marked genes, which had a very narrow range of significantly higher *tau* scores compared to all *Arabidopsis* genes, consistent with PRC2 restricting their expression during development (**Supplemental Fig. S4C**).

To determine which TRGs are actively silenced by H3K9me2 in *Arabidopsis*, we re-analysed gene expression in mutant lines that perturb H3K9me2 function, namely in mutants of the H3K9me2-binding protein AGDP1, which links DNA methylation with H3K9me2, and triple mutants of SUVH4/5/6, the SET domain proteins responsible for H3K9me2 deposition (Zhang et al. 2018). TRGs were significantly enriched among the genes derepressed in *agdp1* and *suvh456* mutants, which was not the case for down-regulated genes (**Supplemental Fig. S4D-E**). This confirmed that H3K9me2 is causal for silencing a subset of 40 TRGs in *Arabidopsis* and indicated that they may be transcribed in particular developmental contexts. Indeed, the majority (87.5%; 35/40) of these *suvh456-*dependent H3K9me2-marked TRGs had strongly enriched expression in reproductive stages, particularly in the pollen vegetative cell nucleus (VN) and developing embryos, with some also expressed in the root (**Supplemental Fig. S4F**). Collectively, these results show that H3K9me is deposited at a substantial number of genes across land plants but is biased towards evolutionarily young TRGs, some of which have been incorporated into gene expression programs that become active mainly during reproductive development in *Arabidopsis*.

In contrast to H3K9me2, genes marked with H3K27me3 in land plants were significantly enriched for those that arose after the emergence of the Streptophyta, whereas species-specific TRGs were significantly depleted for H3K27me3 among the flowering plants (**Fig. 2B-F**). H3K27me3-marked genes were also enriched among genes that emerged in the Bryopsida (or true mosses) in *Physcomitrium* and to *Anthoceros-*specific genes (**Fig. 2B-C**). In the flowering plants *Spirodela*, *Oryza* and *Arabidopsis*, H3K27me3- marked genes were also significantly enriched for genes that arose during the evolution of flowering plants (Magnoliopsida) and their respective phylogenetic orders (**Fig. 2D-F**). Genes in older phylogenetic ranks (i.e., genes that emerged between the first cellular organisms and the Streptophytina) were significantly depleted among H3K9me- and H3K27me3-marked genes (**Fig. 2B-F)**. The unicellular green alga *Chlorella* showed a different pattern compared to the land plants since TRGs were instead significantly enriched among H3K27me3-marked genes (**Fig. 2A**), which incidentally harbours a distinct form of constitutive heterochromatin compared to land plants (**Fig. 1F**). The bias of H3K27me3 at *Chlorella*-specific genes is suggestive of a recent wave of PRC2 recruitment to genes that occurred during evolution of the Trebouxiophyceae.

Gene ontology (GO) enrichment analysis revealed that the *Arabidopsis* genes marked with H3K9me2 or H3K27me3 were enriched for core metabolic processes within the oldest phylogenetic ranks (**Supplemental Fig. S5A; Supplemental Table S4**). H3K27me3-marked genes that emerged in the Streptophyta, Streptophytina and Magnoliopsida (ranks 4, 5 and 6) were additionally enriched for developmental processes, including cell differentiation, tissue and organ development. Developmental processes, together with transcriptional regulation, became the dominant over-represented GO categories among H3K27me3-marked genes that emerged in the Magnoliopsida and Camelineae (ranks 6 and 9). To further probe the bias of PRC2 towards developmental genes, we analysed the partitioning of genes associated with the GO term “development” (GO:0032502) among genes marked with or without H3K27me3 in each phylogenetic rank (**Supplemental Fig. S5B; Supplemental Table S4**). Developmental genes marked with H3K27me3 were already found to be significantly enriched within the oldest phylogenetic rank (Cellular organisms) and included several MIKC-type MADS box TFs such as *AGAMOUS*, 14 *AGAMOUS-LIKE* genes (*AGL*), 4 *SEPALLATA* genes (*SEP1-4*), *PISTILLATA*, *MADS AFFECTING FLOWERING 5* (*MAF5*) and *FLOWERING LOCUS C* (*FLC*) **(Supplemental Fig. S5B; Supplemental Table S4)**. The enrichment of H3K27me3-marked developmental genes became more pronounced in the Viridiplantae, Streptophyta and Magnoliopsida (ranks 3, 4 and 6), with *BLADE-ON- PETIOLE1 (BOP1*), *FLOWERING WAGENINGEN (FWA)* and *PLETHORA 1* (*PLE1*) all notable examples within each rank, respectively. In summary, our analysis has revealed a striking correlation between the evolutionary age of plant genes and the type of heterochromatin used to silence their transcription. Evolutionarily younger genes are more likely to be silenced with heterochromatin marks normally associated with TE silencing, whereas PRC2 silencing of developmental genes has deep roots in the green lineage and became more pronounced upon the advent of complex multicellularity.

## Epigenetic silencing of repeat elements is distinguished by the class of plant TEs

Since distinct forms of heterochromatin preferentially silences genes of differing evolutionary age, we wondered whether this phenomenon would also extend to TEs. Unlike genes however, TEs vary significantly in both sequence and copy number between and even within the same species, making it challenging to assign them to a distinct phylogenetic age. We thus estimated TE age by computing their divergence from the consensus sequence of their assigned family, which serves both as a proxy of evolutionary age and as a prediction of recent transpositional activity (Chalopin et al. 2015). We then compared the relative age and distribution of DNA, LTR and MITE transposons marked with H3K9me, H3K27me3 or both in the six different species.

In *Chlorella*, a large proportion of these TEs (68.8%; 1235/1796) are marked with both H3K9me2 and H3K27me3 (**Fig.1F**; **Fig. 3A**), which are overwhelmingly classified as LTR elements (**Fig. 3B**). The sequence divergence of these co-targeted TEs is significantly lower than that of TEs marked with only H3K9me2 or H3K27me3, indicating that they are relatively young and likely to be recently active in *Chlorella* (**Fig. 3C; Supplemental Fig. S6A**). The second largest proportion of TEs in *Chlorella* are marked solely with H3K27me3 (29.1%; 523/1796), which are significantly enriched for DNA transposons and, given their significantly higher sequence divergence, appear to be oldest group of silenced TEs (**Fig. 3A-C; Supplemental Fig. S6A**). In *Anthoceros*, H3K9me1 and H3K27me3 co-localise at a large distribution of TEs (28.0%; 9182/32741) that are significantly enriched for LTR elements and that are also significantly younger than other silenced TEs (**Fig. 3D-F**). In contrast, the TEs marked solely with either H3K9me1 (57.2%; 18723/32741) or H3K27me3 (14.8%; 4836/32741) were subtly albeit significantly over-represented for either LTR or DNA transposons, respectively, with the latter representing the significantly oldest group (**Fig. 3E-F; Supplemental Fig. S6B**). *Physcomitrium* showed a different pattern since H3K9me2 solely marks the majority of TEs (91.4%; 21158/23138), with only a miniscule proportion (0.19%; 43/23138) being marked with both forms of histone methylation (**Fig. 3G-H; Supplemental Fig. S6C**). Interestingly, the TEs marked solely with H3K27me3 in *Physcomitrium* were also significantly over-represented for DNA transposons and once again significantly older than those marked solely with H3K9me2 (**Fig. 3H-I**).

**Figure 3.**
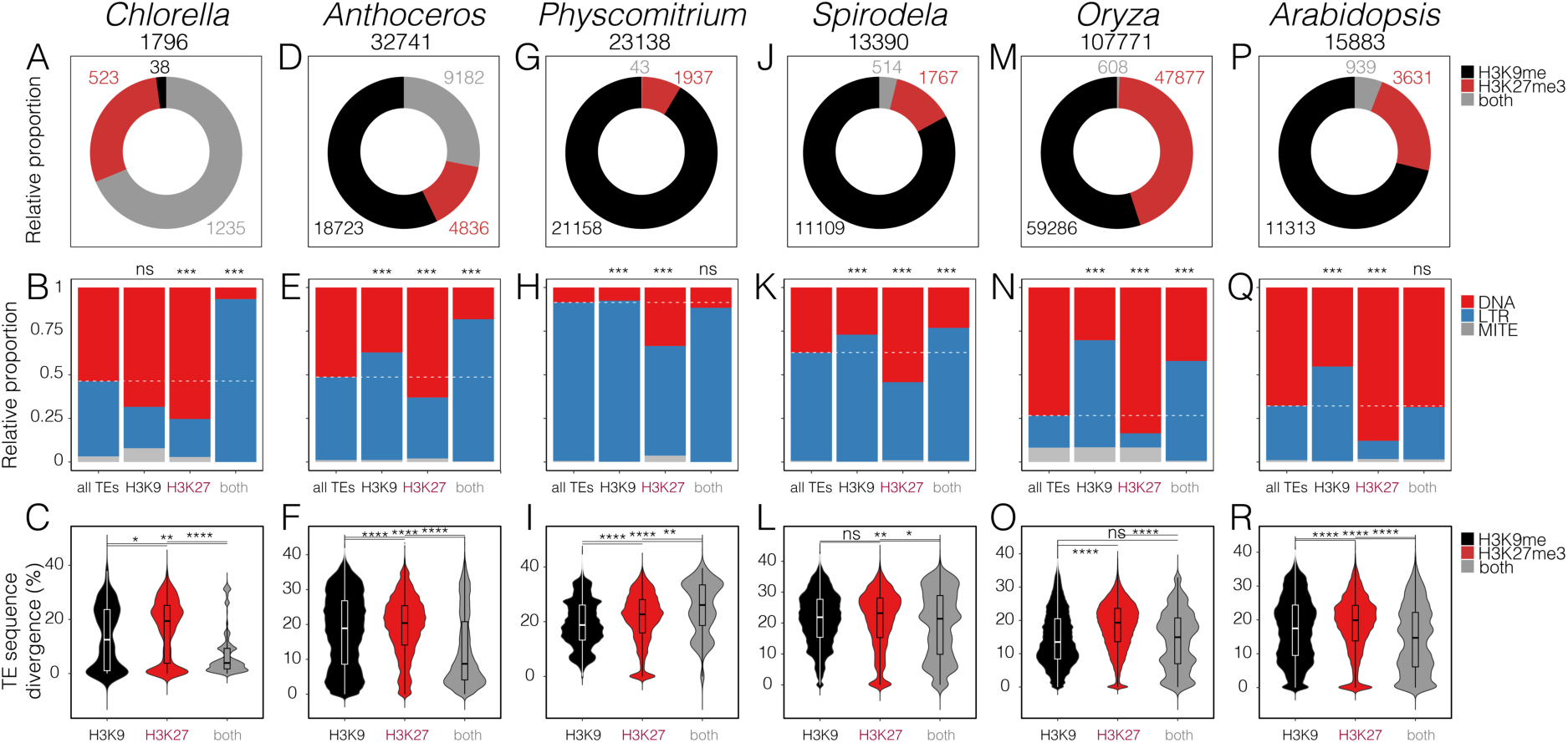
Evolutionary age and epigenetic state of TEs across the green lineage. A trio of panels are grouped for *Chlorella* (**A-C**), *Anthoceros* (**D-F**), *Physcomitrium* (**G-I**), *Spirodela* (**J-L**), *Oryza* (**M-O**) and *Arabidopsis* (**P-R**) under their respective species name. The top element of the trio is a pie chart summarising the relative proportion of DNA, LTR and MITE transposons marked with H3K9me1/2 only (black), H3K27me3 only (red) or both (grey) in each species (A,D,G,J,M,P). The total number of DNA, LTR and MITE TEs present in each species are indicated. The middle element is a stacked barchart summarising the relative proportion of DNA (red), LTR (blue) and MITE (grey) transposons marked with H3K9me1/2 only, H3K27me3 only or both in each species (grouped by column, species indicated above) (B,E,H,K,N,Q). The dashed line marks the relative proportion of DNA and LTR elements for all genes in the genome (i.e., the genome average). Bonferroni corrected *p*-values of a chi-square analysis on top of each bar indicate whether the proportions observed differ significantly from the genome average. The bottom element is a violin plot showing the relative age of TEs marked with H3K9me1/2 only (black), H3K27me3 only (red) or both (grey) in each species (C,F,I,L,O,R). TE age is represented by the sequence divergence of a given TE from the consensus sequence of its assigned family, with higher divergence indicative of an older age and reduced potential for transposition. Each boxplot indicates minimum and maximum values as well as 25^th^, 50th, and 75th quartiles. Significance *p*-values are the result of a Wilcoxon test. * (*p*-value < 0.05), ** (*p*-value < 0.01), *** (*p*-value < 0.001), ns = no significance.

The bias of H3K27me3 towards silencing DNA transposons was also evident among the flowering plants given their significant enrichment among H3K27me3-marked TEs in *Spirodela* (**Fig. 3J-K**), *Oryza* (**Fig. 3M-N**) and *Arabidopsis* (**Fig. 3P-Q**). Conversely, the TEs solely marked with H3K9me2 in flowering plants were significantly enriched for LTR elements (**Fig. 3K,N,Q**). The TEs marked solely with H3K27me3 in *Oryza* and *Arabidopsis* also had significantly higher levels of sequence divergence compared to those marked solely with H3K9me2 (**Fig. 3O,R; Supplemental Fig. S6E-F**), suggesting that H3K27me3 is preferentially found at ancient domesticated TEs, as has been reported in *Arabidopsis* (Hisanaga, Romani, et al. 2023; Hure et al. 2025). Moreover, unlike in *Chlorella* and *Anthoceros*, H3K9me and H3K27me3 were co-deposited at very few TEs in *Physcomitrium* and flowering plants, consistent with the functional bifurcation of these epigenetic pathways during land plant evolution (**Fig. 3A,D,G,J,M,P**). Our analysis has thus revealed that, unlike for genes, the evolutionary age of TEs does not correlate with distinct forms of heterochromatin. Instead, the epigenetic silencing of TEs appears to be influenced by the mode of TE transposition, particularly among land plants, where H3K9me2 and H3K27me3 preferentially mark LTR and DNA transposons, respectively.

## Neighbouring transposons spread heterochromatin to evolutionarily young genes

Heterochromatic H3K9me domains typically span across large chromosomal domains to promote silencing of repetitive DNA and TEs, which can sometimes spread beyond their intended targets and repress neighbouring genes (Cutter DiPiazza et al. 2021). To assess whether the preference of H3K9me deposition for TRGs might be caused by heterochromatin spreading, we compared the chromosomal distribution of the phylogenetically-ranked genes in each species (**Fig. 4**). In *Anthoceros*, the density of young genes was prominent within regions with a higher density of TEs and H3K9me1 peaks (**Fig. 4A**), which was further reflected in metaplots of relative TE density (**Fig. 4B**). Consistently, *Anthoceros*-specific genes (rank 6) were the only group to be significantly over-represented for genes lying in the vicinity of H3K9me1-marked TEs (**Fig. 4B-C; Supplemental Fig. S7A-B**). Although these observations were less obvious in *Physcomitrium*, TRGs (rank 7) were nevertheless also significantly enriched for genes within 1kb of a TE and/or H3K9me2 peak (**Fig. 4D-F; Supplemental Fig. S7C-D**). Thus, TRGs are significantly more likely to lie adjacent to heterochromatic TEs in both bryophyte species, explaining their increased propensity for silencing with H3K9me1/2 (**Fig. 2C-F**).

**Figure 4.**
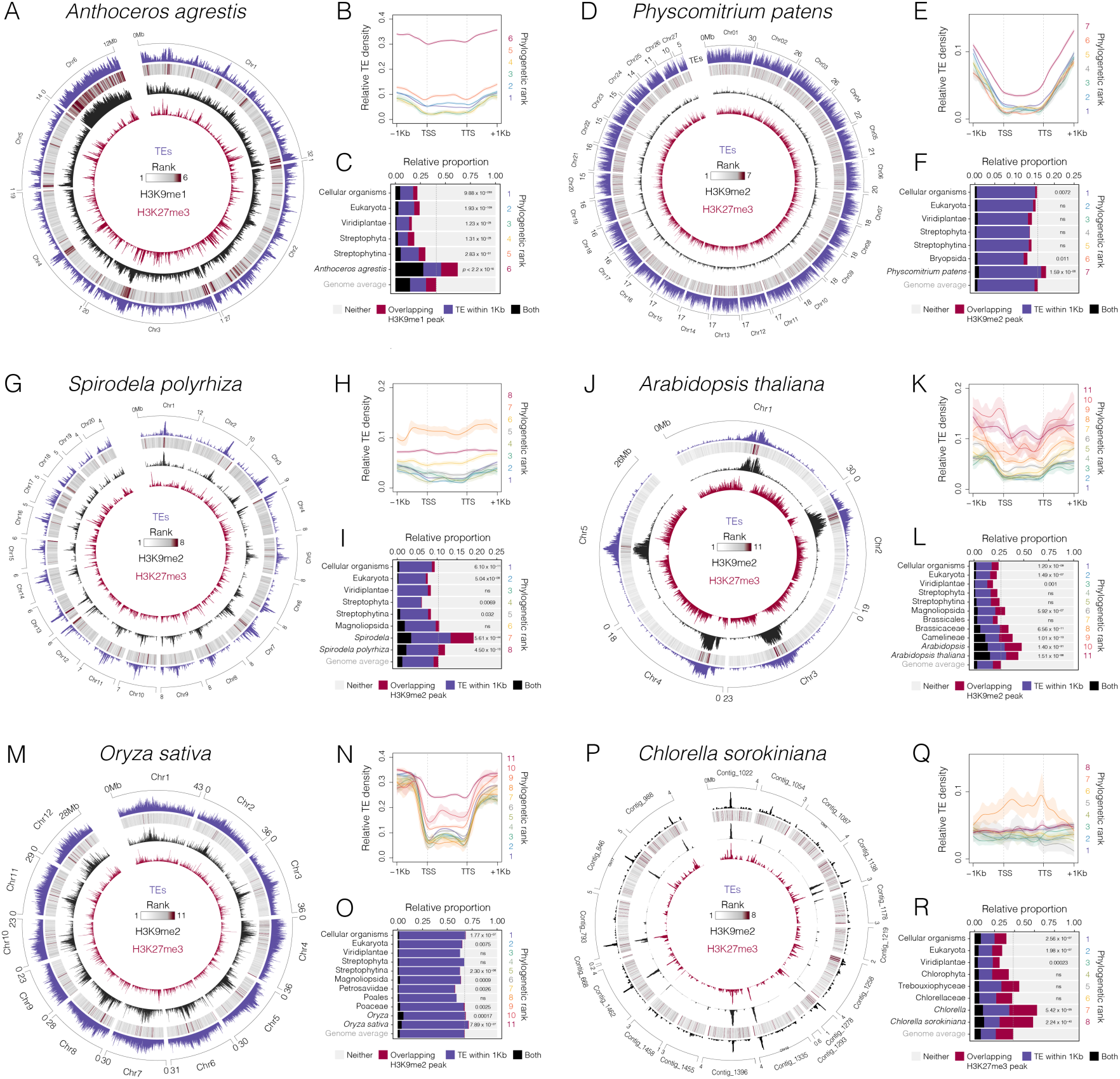
Neighbouring TEs spread constitutive heterochromatin to evolutionarily young genes. A trio of panels are grouped for *Anthoceros* (**A-C**), *Physcomitrium* (**D-F**), *Spirodela* (**G-I**), *Arabidopsis* (**J-L**), *Oryza* (**M-O**) and *Chlorella* (**P-R**) under their respective species name. The first element of the trio24rabidcircos plot (A,D,G,J,M,P) showing the chromosomal density (in 10kb bins) of repeats and TEs (purple), average phylogenetic rank of genes (grey-claret heatmap), H3K9me2 peaks (H3K9me1 for *Anthoceros*) (black) and H3K27me3 peaks (claret). The second element is a profile plot (B,E,H,K,N,Q) of relative TE density centred on each group of phylogenetically-ranked genes, where shading indicates the standard error of TE density. The third element is a barplot showing the relative proportion of genes in each phylogenetic rank that lie within 1 kb of a TE (purple), that overlap with an H3K9me1/2 peak (black) (or H3K27me3 peak in *Chlorella* in green), or both (claret). The dashed line marks the maximum proportion of these three combined categories for all genes in the genome (i.e., the genome average). Bonferroni corrected *p*-values of a chi-square analysis indicate whether the proportions observed in each phylogenetic rank differ significantly from the genome average.

In *Spirodela* and *Arabidopsis*, TEs are known to be enriched within pericentromeric chromosomal regions that also happen to be relatively depleted of genes (Kaul et al. 2000; Dombey et al. 2024). The density of TRGs was increased within TE-rich pericentromeric regions in both *Spirodela* (ranks 7-8) and *Arabidopsis* (ranks 8-11) (**Fig. 4G,J**). Consistently, the relative proportion of young genes lying in close proximity of TEs and H3K9me2 peaks was significantly increased when compared with older genes, which was further reflected in metaplots of TE density (**Fig. 4H-I,K-L; Supplemental Fig. S7E-H**). Although the pericentromeric regions of the *Oryza* genome are less obvious than in *Spirodela* and *Arabidopsis*, they nonetheless manifest as distinct chromosomal regions that are relatively enriched with TEs (**Fig. 4M**). These TE-enriched hotspots were again found to have a higher density of young genes (ranks 9-11) compared to neighbouring chromosomal regions (**Fig. 4M**). Although more subtle when compared to other flowering plants genomes, TRGs in *Oryza* were once again significantly more associated with H3K9me2-marked TEs and had more TE insertions within their gene body compared with older genes (**Fig. 4N-O; Supplemental Fig. S7I-J**). Our results thus demonstrate that TRGs are more commonly found within heterochromatic TE-rich regions of flowering plant genomes, which likely results in their silencing by the spreading of H3K9me2 domains from neighbouring TEs.

Despite the obvious TE-rich hotpots present in the *Chlorella* genome, we did not observe an increased density of young genes as seen within pericentromeric regions of the flowering plant genomes (**Fig. 4P**). Nevertheless, TRGs in *Chlorella* (ranks 7-8) were found significantly more frequently in the vicinity of H3K27me3-marked TEs (**Fig. 4Q-R; Supplemental Fig. S7K**). Interestingly, most TRGs in *Chlorella* that overlapped with an H3K27me3 domain had no adjacent TEs within 1 kb (**Fig. 4R; Supplemental Fig. S7L**), suggesting that PRC2 recruitment of TRGs is not simply caused by H3K27me3 spreading from neighbouring TEs but might rather involve a TE-independent PRC2 targeting mechanism.

## PRC2 regulates a conserved group of gene families across the green lineage

Our epigenetic analysis of gene ages has revealed that H3K27me3 preferentially marks genes that emerged during and after Streptophyte evolution. PRC2 thus appears to have been recruited to a swathe of genes during this evolutionary period, suggesting that modern-day land plants may share a network of PRC2- regulated gene families. To investigate this, we performed an orthology inference analysis to group gene families from the six representative species into 18,415 orthogroups, then calculated the relative proportion of genes marked with H3K27me3 (**Supplemental Table S5; Supplemental Table S6**). Hierarchical clustering of the 1,520 orthogroups that were deeply-conserved in all six species revealed that those regulated by PRC2 are largely unique to each species, consistent with the distinct evolutionary history of each organism (**Fig. 5A**). A Cochran’s Q test (Q = 4276.5, *p* < 2.2 x 10^-16^) confirmed that the observed proportions of H3K27me3-marked orthogroups across the six species was not random. Further analysis revealed a number of orthogroups that were commonly marked with H3K27me3 between different species, including a core of 92 gene families (6.1%; 92 of 1520) common to the five land plants we analysed (**Fig. 5B**), a proportion that significantly exceeded the null expectation of 0.15% (*p* < 2.2 x 10^-16^, binomial test).

**Figure 5.**
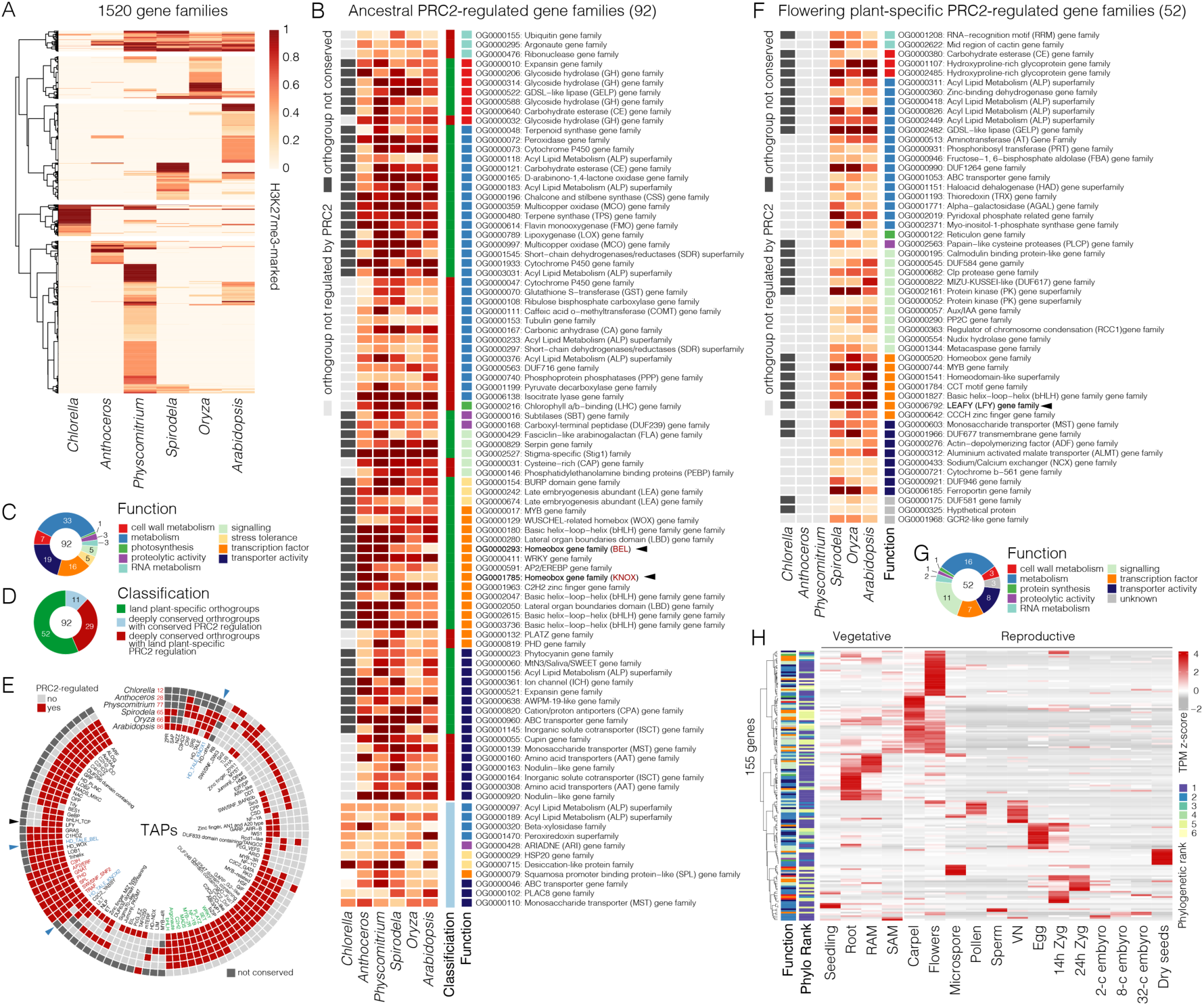
The conserved topology of PRC2-repressed gene networks across land plants. (**A**) Heatmap summarising the 1520 orthogroups (or gene families) conserved across the six species analysed in this study. The relative proportion of genes in each orthogroup marked with H3K27me3 in each species are indicated and coloured according to the inset scale. (**B**) Heatmap showing the cluster of gene families marked with H3K27me3 in all five land plants using the colour scale in panel B. Conserved orthogroups containing no genes marked with H3K27me3 in *Chlorella* are shown in light grey, whereas orthogroups not conserved in *Chlorella* are shown in dark grey (see classification column and panel D). Assigned functions are coloured according to panel C. (**C**) Pie chart summarising the different functions of the gene families in panel B. (**D**) Pie chart summarising the number of orthogroups in panel B. This was composed of PRC2-regulated gene families only conserved in the five land plants (green), gene families conserved in *Chlorella* but only marked with H3K27me3 in land plants (red), and PRC2-regulated gene families deeply-conserved in all six species (light blue). (**E**) Circular heatmap summarising the 102 transcription-associated protein (TAP) families regulated by PRC2 across each species. TAPs with deeply- conserved regulation are indicated in red, land plant-specific regulation in green, and BELL-KNOX homologs in blue. (**F**) Heatmap showing PRC2-regulated gene families specific to flowering plants. (**G**) Pie chart summarising the different functions of the gene families in panel F. (**H**) Heatmap illustrating the developmentally regulated expression of the *Arabidopsis* PRC2-regulated gene families specific to flowering plants in panel F. Gene functions are coloured according to the annotations in panel G, while the phylogenetic rank of genes is coloured according to the scale to the right of the heatmap. Expression represents the z-score of normalised RNA-seq TPM values.

We divided this core set of PRC2-regulated orthogroups into three main groups and further classified their biological function based on gene annotations in *Arabidopsis* (**Fig. 5C-D; Supplemental Table S7**). The largest group of gene families encoded catabolic enzymes involved in various metabolic functions, including lipid biogenesis, cell wall metabolism and proteins that modulate responses to osmotic and oxidative stress (**Fig. 5B-C**). Some signalling processes were also present and included a Phosphatidylethanolamine binding protein (PEBP) gene family related to *FLOWERING LOCUS T* and *TERMINAL FLOWER 1* (Wickland and Hanzawa 2015). Gene families involved in transport activity of inorganic ions, sugars and hormones were also abundant, as were a defined set of TF families (**Fig. 5B-C**). While the majority of PRC2-regulated gene families were only conserved in the five land plants (52 of 92 orthogroups; 56.0 %), around a third were also conserved in *Chlorella* but only marked with H3K27me3 in land plants (29 of 92 orthogroups; 31.5 %), suggesting that PRC2 was recruited to these gene families upon or after the emergence of the Streptophyta (**Fig. 5D**). The final group was composed of a small subset of gene families marked with H3K27me3 in all six species (11 of 92 orthogroups; 0.5%), highlighting a core of deeply-conserved PRC2-regulated gene families involved in core biological processes, including ion and sugar transport, lipid and carbohydrate metabolism, and stress tolerance (**Fig. 5B-D; Supplemental Table S8)**. This deeply-conserved group of PRC2 targets also included Squamosa promoter binding protein−like (SPL) genes (**Fig. 5B**), which belong to an ancient family of plant-specific TFs that play diverse roles in land plants (Preston and Hileman 2013). Our analysis has thus revealed how PRC2 regulates a core of orthologous genes across land plants as well as in their unicellular green algal relatives.

Among the 103 transcription-associated proteins (TAPs) regulated by PRC2 across the six species, only 16 were marked with H3K27me3 in all five land plants, which included members of the AP2/ERF, bHLH, C2H2, R2R3-MYB and WRKY family of TFs (**Fig. 5E**; **Supplemental Table S9**). Argonaute and SET domain proteins were also marked with H3K27me3 in the land plants, highlighting how PRC2 targeting also evolved to silence major components involved in small RNA and chromatin regulation (**Fig. 5E**; **Supplemental Table S9**). Although only thirteen TAPs were marked with H3K27me3 in *Chlorella*, seven of these were shared with the five land plants and included AP2/ERF, PHD and SPL type TFs and SWI/SNF chromatin remodelling proteins (**Fig. 5E**). Most notable among the TFs with conserved PRC2 regulation were two orthogroups descended from the TALE-HD TFs KNOX (KNOTTED-like homeobox) and BELL (BEL-Like) (**Fig. 5B,E**). KNOX/BELL have deep evolutionary origins in unicellular green algae (Lee et al. 2008; Nishimura et al. 2012; Kariyawasam et al. 2019) and play a conserved role in sexual reproduction by activating the zygotic program after fertilisation in *Chlamydomonas* and *Marchantia* (Sakakibara et al. 2013; Horst et al. 2016; Dierschke et al. 2021; Hisanaga et al. 2021). Although we were unable to identify any obvious homologs of KNOX/BELL in *Chlorella*, which is assumed to reproduce asexually (Hovde et al. 2018), the KNOX/BELL orthologs present in *Anthoceros* and *Physcomitirium* were all marked with H3K27me3 (**Fig. 5B; Supplemental Fig. S8**). Analysis of published CUT&RUN data in *Marchantia* also confirmed that most KNOX/BELL orthologs are also silenced with H3K27me3 in liverworts (**Supplemental Fig. S8**), which has been demonstrated functionally in previous work (Hisanaga, Romani, et al. 2023). As the KNOX/BELL gene families expanded during flowering plant evolution, PRC2 regulation appears to have also diversified by targeting a subset of KNOX/BELL homologs (**Fig. 5B**). Other noteworthy examples of TF orthogroups with deeply conserved PRC2 regulation included members of the WOX, WRKY, bHLH and DREB-A2 subfamily of AP2/ERF TFs, which regulate a wide range of vital processes in *Arabidopsis* ranging from seed dormancy, embryogenesis, stomatal patterning through to abiotic stress responses (**Fig. 5B**). Key transcriptional regulators of the plant life cycle and cellular patterning thus appear to be highly conserved target genes of PRC2 in land plants.

Interestingly, the TAP families regulated by PRC2 in flowering plants were a larger and more diverse group compared with those in *Chlorella* and *Anthoceros* (**Fig. 5E**), emphasising the broader control PRC2 evolved to exert over gene regulatory networks in flowering plants. *Physcomitrium* was an exception since the number of H3K27me3-marked TAPs were similar to those in flowering plants, although at least seven were exclusive to this species, including E2F/DP and HMG-type TFs (**Fig. 5E**). The expansion of PRC2-regulated TAPs among the flowering plants raised the question as to whether they also share a distinct network of PRC2-regulated gene families. Within the H3K27me3-marked orthogroups were a group of 52 gene families (3.4%; 52 of 1520) that were specifically regulated by PRC2 in *Spirodela*, *Oryza* and *Arabidopsis* but not in bryophytes, a proportion that significantly exceeded the random expectation of 2.3% (*p* < 0.036, binomial test) (**Fig. 5F**). While many of the functions were broadly similar to those observed in the PRC2-regulated gene network common to the land plants, the flowering plant-specific network was notably more diverse in signalling processes (**Fig. 5F-G**). This included gene families encoding two distinct types of protein kinases, a group of ABA-induced Type 2C protein phosphatases (PP2C) as well as AUX/IAA genes involved in auxin responses (**Fig. 5F-G**). Several TF families were also present and included the floral meristem identity regulator LEAFY and a CCT MOTIF FAMILY (CMF) group of TFs related to ASML2, a TF with high expression in reproductive organs that regulates sugar-inducible genes (**Fig. 5E-F**) (Weigel et al. 1992; Masaki et al. 2005). Transcriptomic analysis of the flowering plant-specific PRC2 network of genes in *Arabidopsis* revealed strong developmentally-regulated expression in reproductive and vegetative organs and tissues that distinguish the flowering plant lineage, including flowers, pollen, seeds and roots (**Fig. 5H**). These data highlight how PRC2 regulates a conserved network of genes specifically among flowering plants, which includes key transcriptional regulators and signalling proteins that specify major organs and tissues characteristic of this diverse group.

## H3K27me3 domains in land plants share common *cis*-regulatory motifs

In animals and plants, H3K27me3 deposition is targeted to specific loci by TFs that tether PRC2 to *cis*- regulatory regions called Polycomb response elements (PREs) (Blackledge and Klose 2021). We thus pondered whether the type of TFs that recruit PRC2 to H3K7me3-marked genes might be similar among the six species. To address this, we performed motif enrichment analysis to determine which TFs are likely to bind within the H3K27me3 domains in each species. Because the binding specificity of orthologous TFs is known to be highly conserved deep across evolutionary time (Nitta et al. 2015), we reasoned that the binding preference of *Arabidopsis* TFs could be used to identify potential PRE elements. For this we used a set of experimentally-derived DNA binding motifs for *Arabidopsis* TFs generated with DAP-seq (O’Malley et al. 2016), but focused only on DNA binding motifs from TF families that had at least one ortholog present in all five species.

In total we identified 234 significantly enriched motifs within H3K27me3 domains across all six species (**Fig. 6A; Supplemental Table S10**). Among these were 8 motifs that were commonly enriched across all five land plants, with the majority (87.5%; 7/8) bound by three distinct families of AP2/ERF TFs (**Fig. 6A-C; Supplemental Table S10**). In contrast, non-H3K27me3-marked genes that are also conserved across the five species were not enriched for the same motifs (**Supplemental Table S10**). H3K27me3 domains in *Chlorella* also showed no enrichment for AP2/ERF motifs, despite the presence of AP2/ERF TFs in *Chlorella* (**Fig. 5E**; **Supplemental Table S9**). To further probe the potential relationship of these TFs in PRC2 recruitment, we plotted the occurrence of the commonly enriched motifs over PRC2 target genes alongside the profile of H3K27me3 (**Fig. 6C-E**). In each of the five land plants, these motifs occurred in a pattern that mirrored the profile of H3K27me3 (**Fig. 6E**). Although some motifs were also present in the upstream flanking regions of the genes, most were positioned downstream of the TSS along the gene body (**Fig. 6E**). This is consistent with recent reports of several plant TFs binding within the body of genes, including PRE-binding TFs in *Arabidopsis* (Xiao et al. 2017; Voichek et al. 2024).

**Figure 6.**
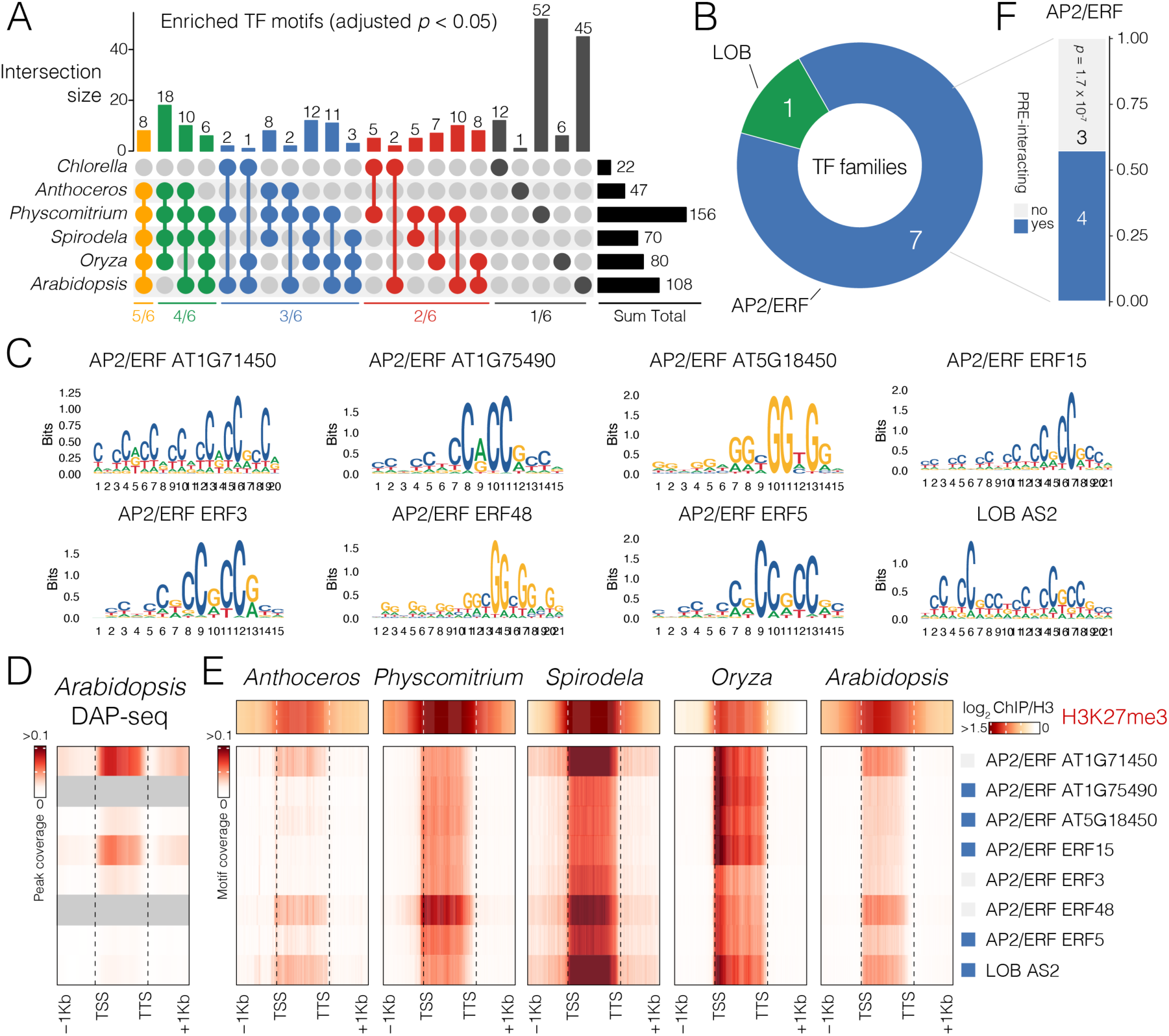
H3K27me3 domains share DNA-binding motifs across large evolutionary distances. (**A**) Upset plot summarising the overlap and number of DNA binding motifs significantly enriched (adjusted *p*-value < 0.05) within H3K27me3 domains in each land plant species (see Fig. 4B; Supplemental Table S7). Yellow: enriched in five of six species, Green: four of six, Blue: three of six, Red: two of six, Gray: enriched in only one species. (**B**) The number and type of DNA binding motifs that are commonly enriched across all land plants in panel A (yellow bar). (**C)** PWM logos of the commonly enriched DAP-seq motifs in panel B.(**D**) Averaged signal of DAP-seq peaks centred on H3K27me3-marked genes in *Arabidopsis*. (**E**) Occurrence of the commonly enriched DNA binding motifs (panel B) centred on H3K27me3-marked genes in each land plant species. ChIP-seq log_2_ ratio of H3K27me3 enrichment relative to H3 over the same genes is indicated in the heatmaps above. Blue boxes indicate PRE- binding TFs in *Arabidopsis* (Lodha et al. 2013; Xiao et al. 2017). (**F**) Proportion of the AP2/ERFs in panel B that are known to bind to PREs in *Arabidopsis.* (Lodha et al. 2013; Xiao et al. 2017). Significance *p*-value is the result of a Fisher’s exact test based on the overlap between the AP2/ERF TFs with a commonly enriched motif and the 233 PRE- binding TFs identified in Xiao *et al*., 2017.

Because the DAP-seq dataset assayed TF binding to naked *Arabidopsis* genomic DNA, DAP-seq peaks can give an accurate representation of *in vivo* TF binding potential. Where available, the DAP-seq peaks of TF binding were also enriched over gene bodies, particularly for AP2/ERF TFs, demonstrating that the pattern of motif predictions mirrors TF binding in a genomic context (**Fig. 6D**). Interestingly, AP2/ERF TFs are among the most common family of TFs that can bind to PRE-like sequences in *Arabidopsis,* many of which can also interact with PRC2 (Xiao et al. 2017; Xie et al. 2022). Four of the seven AP2/ERF TFs we identified have been shown to physically bind PREs in *Arabidopsis*, an association that is statistically supported (**Fig. 6F**). Motifs of the LOB domain TF ASYMMETRIC LEAVES 2 (AS2) were also commonly enriched, which is known to physically recruit PRC2 to its target loci in *Arabidopsis* (**Fig. 6B-C,E**) (Lodha et al. 2013). Thus, H3K27me3 domains share similar DNA binding sites across distantly-related land plants, highlighting a conserved and potentially ancestral feature of PRC2-regulated genes in plants.

## Discussion

The genomic era has heralded essential insights into the genetic bases of land plant evolution and points to the expansion and diversification of a genetic toolkit with deep origins in their aquatic algal ancestors (Bowman 2022; Dhabalia Ashok et al. 2024). This includes most of the TF families found in modern-day plants, many of which play conserved roles in environmental and developmental regulation across large evolutionary distances (Romani and Moreno 2021). Although these TFs would have undoubtedly shaped gene regulatory networks in the last common Streptophyte ancestor, added layers of gene regulation would have been necessary to modulate TF expression and define new cell types and developmental traits. Here, we have described the evolutionary dynamics of epigenetic regulation across the Viridiplantae and show how epigenetic regulatory networks have progressively expanded and diversified during land plant evolution.

Our analysis has revealed that evolutionarily young or TRGs are highly over-represented among genes that are silenced by constitutive heterochromatin in plants (**Fig. 4**). Closer inspection of the chromosomal distribution of genes grouped by their evolutionary age revealed that evolutionarily older genes are depleted from heterochromatic regions like pericentromeres. In contrast, TRGs were enriched in TE-rich heterochromatic regions of several plant species, increasing their likelihood for silencing with constitutive heterochromatin. In *Drosophila*, TRGs tend to emerge within pre-existing repressive domains that are enriched with H3K27me3 (Zhang and Zhou 2019). In the nematode *Pristionchus pacificus*, TRGs are also strongly enriched for heterochromatic states marked with H3K27me3, H3K9me3 or both (Werner et al. 2018). In the brown alga *Ectocarpus*, TRGs are highly enriched along UV sex chromosomes that are distinguished by a brown algal-specific repressive chromatin state enriched with H3K79me2 (Gueno et al. 2013; Luthringer et al. 2015). TRGs thus appear to be partitioned into repressive chromatin domains in a diverse range of eukaryotes and tend to be transcribed in a restricted number of cell types and tissues (Luthringer et al. 2015; Werner et al. 2018; Zhang and Zhou 2019). Our analysis has revealed a similar pattern in *Arabidopsis* where some Brassicaceae-specific H3K9me2-regulated genes are preferentially transcribed in reproductive cell types but also in the root (**Supplemental Fig. S4**). The expression of these H3K9me2-regulated TRGs in pollen, embryos and roots is consistent with the reprogramming of constitutive heterochromatin reported in these developmental stages (Schoft et al. 2009; Kawakatsu et al. 2016; Papareddy et al. 2020; Borg, Papareddy, et al. 2021; Parent et al. 2021). Increasing evidence across eukaryotes suggests that TRGs can play key biological roles, including in *Arabidopsis* (Fakhar et al. 2023), highlighting these H3K9me2-regulated TRGs as potentially interesting candidates for future functional characterisation. Interestingly, pollen is a known hotspot for new gene emergence in flowering plants (Wu et al. 2014), suggesting that the integration of TRGs into the H3K9me2 silencing pathway could be a potential route for the evolution of tissue-specific expression and the selection of novel gene functions. Thus, although PRC2 appears to dominate gene silencing across land plants, lineage-specific cases of H3K9me2-regulated gene networks appear to have emerged during flowering plant evolution.

Our analysis of chromatin in the unicellular green microalga *Chlorella sorokiniana* suggests that two types of heterochromatin mediate silencing in *Chlorella*, which are both composed of H3K27me3 but differentiated by the presence or absence of H3K9me2 (**Fig. 1)**. Our analysis also revealed a substantial proportion of genes marked with H3K27me3, indicating that PRC2 regulation of genes has deep origins in the green lineage. H3K9me2 and H3K27me3 co-occur together at most TEs in *Chlorella*, which is strikingly reminiscent of their co-deposition at TEs in *Paramecium* (Frapporti et al. 2019). This indicates a common evolutionary history shared by these two epigenetic pathways in unicellular eukaryotes prior to their functional bifurcation later during evolution. In contrast, H3K27me3-marked genes in *Chlorella* were devoid of H3K9me2 and generally did not lie in the vicinity of LTR or DNA transposons, highlighting a separation between the chromatin states that define TEs and genes in this microalga. This suggests that *cis* recruitment of PRC2 to genes in *Chlorella* occurs by an unknown mechanism, potentially via PRE-like elements commonly found in more complex multicellular eukaryotes (Blackledge and Klose 2021). Unicellular eukaryotes like green microalgae could thus make interesting model systems to study the origin and dynamics of PRC2 recruitment in the future.

By stratifying genes by their evolutionary age and epigenetic state in multiple species of land plants, we observed that genes specific to the Streptophytina lineage are significantly over-represented among the genes targeted by PRC2 (**Fig. 2**), which coincides with the point in evolution when multicellularity first arose in the charophyceaen macroalgae (Umen 2014). A significant over- representation of PRC2-target genes is also evident among genes that emerged during the rise of the embryophytes and subsequent evolution of the monocots and dicots. This suggests a major period of regulatory adaptation during Viridiplantae evolution where hundreds of genes were incorporated into epigenetic networks controlled by PRC2. These landmark events are associated with considerable *de novo* gene birth and gene family diversification (Bowles et al. 2020; Barrera-Redondo et al. 2023), which may have provided ripe context for new PRC2 gene regulatory networks to emerge and fuel evolutionary change. Based on our knowledge of PRC2 function in the modern-day, this is likely to have been driven by genetic changes through the emergence of motifs bound by TFs that recruit PRC2, so-called PRE-binding TFs (Xiao et al. 2017; Blackledge and Klose 2021). Recent studies have suggested that H3K27me3-marked TE fragments, particularly those originating from DNA transposons, were co-opted as PRE elements during Archaeplastida evolution (Hisanaga, Romani, et al. 2023). Our analysis of the epigenetic state and evolutionary age of TEs suggests that DNA transposons are the preferred target of H3K27me3 across distantly-related Viridiplantae species, including unicellular green microalgae, suggesting an ancient evolutionary relationship (**Fig. 3)**. Our work thus provides further credence to the notion that DNA transposons were domesticated to impact gene regulation during Archaeplastida evolution, and we hypothesise that embryophyte evolution in particular was an important period of epigenetic adaptation given the preferential targeting of PRC2 to genes that emerged during this period of plant evolution.

This raised the question as to whether the functional topology of the gene networks regulated by PRC2 are common or unique among distantly-related plants. Overall, PRC2-regulated gene families are largely unique to the species we analysed, which is unsurprising given the distinct evolutionary history of each lineage (**Fig. 5A**). Nevertheless, our analysis has revealed that PRC2 silences a common network of genes in the vegetative phase of distantly-related land plants (**Fig. 5B**). We also identified an additional PRC2-repressed network shared among flowering plants, which becomes predominantly active during reproductive development in *Arabidopsis* (**Fig. 5F-H**). This could represent an ancestral signature of the PRC2-regulated gene networks established during the evolution of land and flowering plants. Alternatively, these shared networks may have arisen independently through convergent evolution, highlighting the type of genes PRC2 has been honed to silence over the course of evolution. Several glycosyl hydrolase gene families stood out in these networks, which are crucial for carbohydrate and glycoconjugate metabolism during various cellular processes, including the mobilisation of starch reserves, pathogen defence and the assembly and disassembly of the cell wall (Kfoury et al. 2024). In addition to gene families with diverse metabolic functions, several others involved in sugar, amino acid and organic ion transport also form part of these networks, whose fine-tuned epigenetic regulation may facilitate nutrient relocation within specific tissues and organs (Martinoia et al. 2000; Nagata et al. 2008). Interestingly, phytohormone signalling pathways were not part of the PRC2-regulated network shared among the five land plants, despite the fact that auxin signalling pathways are known to be prominent targets of PRC2 in flowering plants (Lafos et al. 2011; Wójcikowska et al. 2020; Wu et al. 2023). Our results show that PRC2 regulation of auxin signalling, specifically AUX/IAA proteins, emerged later during flowering plant evolution, likely in response to the expansion of the minimal auxin signalling systems seen in bryophytes (**Fig. 5F**) (Flores-Sandoval et al. 2015; Suzuki et al. 2023).

The most prominent and well-studied PRC2 target genes in complex multicellular eukaryotes tend to encode for TFs. Our analysis has confirmed this in several land plants but has provided further insight into breadth of TAPs regulated by PRC2 across the green lineage (**Fig. 5E**). The number and diversity of TAPs regulated by PRC2 was markedly increased in the vegetative phase of flowering plants compared to bryophytes, which included not only DNA binding TFs but also key players of small RNA silencing (Argonautes), histone methylation (SET domain proteins) and chromatin remodelling (SWI/SNF proteins). The progressive increase in PRC2-regulated TAP genes was likely key in establishing novel regulatory networks during land plant evolution, potentially facilitating the emergence and specification of new cell types and tissues. Whether the diversity of PRC2-regulated TAPs changes across the complex life cycle of these organisms remains unclear, highlighting an intriguing question to address once more comprehensive datasets become available. Interestingly, the number of TAP families regulated by PRC2 in *Chlorella* was fairly low in number compared to the land plants, despite the fact that many of the same plant TAPs have orthologs in *Chlorella* (**Fig. 5E**). We identified at least seven TAP families with deeply-rooted PRC2 regulation across the green lineage, including AP2/ERF, PHD and SPL type TFs as well as SWI/SNF chromatin remodellers, unveiling an ancient regulatory relationship that has persisted from green microalgae into modern-day bryophytes and flowering plants. Our results also indicate that the master regulator of floral meristem identity LEAFY came under control of PRC2 during flowering plant evolution and forms part of a PRC2-repressed network associated with reproduction in this dominant group (**Fig. 5E,F)**. Interestingly, our analysis of H3K27me3 domains across land plants has revealed a tendency for enrichment with GC-rich TF-binding motifs, which also happen to occur in a pattern that mirrors H3K27me3 deposition (**Fig. 6**). This is reminiscent of mammalian systems where PRC2 is recruited to GC-rich regions by DNA binding proteins that have an affinity for low complexity GC-rich motifs or CpG dinucleotides (Mendenhall et al. 2010; Wachter et al. 2014; Laugesen et al. 2019). The GC-rich motifs we identified are bound by AP2/ERF family TFs and the LOB family TF AS2, which can physically interact with PRC2 in *Arabidopsis* and recruit it to its target loci in a manner similar to that reported in animals (Lodha et al. 2013; Xiao et al. 2017). Because these motifs were not enriched in the H3K27me3 domains of *Chlorella*, it appears that this feature may have arisen later in plant evolution, which could have potentially facilitated the targeting of PRC2 to genes. However, future validation of PRC2-TF interactions in green microalgae and bryophytes will be crucial to clarify this and shed more light on this ancestral regulatory relationship.

Among the shared PRC2-regulated TF families we identified were the heterodimerisation partners KNOX and BELL (Bellaoui et al. 2001). Given the ancestral role these TALE-HD TFs play in activating the diploid zygotic/sporophytic program in *Chlamydomonas reinhardtii* and *Marchantia polymorpha*, it is hypothesised that they were key genetic determinants that sparked the evolution of haploid-diploid life cycles in plants (Lee et al. 2008; Sakakibara et al. 2013; Hisanaga et al. 2021). Our analysis shows that BELL and KNOX are both regulated by PRC2 in the haploid gametophyte of all three major bryophyte groups, suggesting that their regulation by PRC2 may be an ancestral feature of land plants. This regulation is likely to be highly relevant since *KNOX* and *BELL* are derepressed in PRC2 mutants of *Physcomitirium* and *Marchantia*, resulting in the gametophyte displaying sporophyte-like features or lethality, respectively (Pereman et al. 2016; Hisanaga, Romani, et al. 2023). Transcriptional silencing of BELL and KNOX by PRC2 in the haploid phase of the life cycle would have been key for repressing diploid identity, further fuelling the idea that their upstream regulation could have facilitated the evolution of haploid-diploid transitions (Vigneau and Borg 2021). KNOX/BELL homologs appear to have been lost in *Chlorella* so we were unable to assess whether they are also regulated by PRC2. Broader profiling of chromatin landscapes in other green algal lineages and land plants, including the more complex charophycean macroalgae and ferns, is required to better understand when PRC2 silencing of KNOX/BELL first emerged, and also help trace when and how PRC2-regulated gene networks were shaped during the earliest phases of Viridiplantae evolution.

## Materials and Methods

### *Chlorella* ChIP-seq profiling

The unicellular green alga *Chlorella sorokiniana* CCALA 259 (equivalent to UTEX 1230) was obtained from the Culture Collection of Autotrophic Organisms at the Institute of Botany of the Czech Academy of Sciences (Trebon, Czech Republic). The cultures were grown on ½ strength Šetlík-Sluková (SS) medium (Hlavová et al. 2016). For routine subculture, cultures were streaked every three weeks on ½ SS medium solidified with agar (1.5 %, w/v) and grown at an incident light intensity of 100 µmol.m^-2^.s^-1^ of photosynthetically active radiation (PAR). For the experiment, 300 ml of the liquid ½ SS medium was inoculated directly from the plates and the cultures were placed in glass cylinders (inner diameter 30 mm, height 500 mm) at 30 °C and “aerated” with a mixture of air and CO2 (2%, v/v) at a flow rate of 15 l.h^-1^. The cylinders were illuminated from one side with dimmable fluorescent lamps (OSRAM DULUX L55W/950 Daylight, Milano, Italy), the light intensity of which was adjusted so that 500 µmol.m^-2^.s^-1^ of PAR hit the surface of the cylinders. The cultures were grown under continuous light until a cell density of approximately 1×10^8^ cells.ml^-1^ was reached. They were then shifted to 40 °C and a light intensity of 700 µmol.m^-2^.s^-1^ for 24 hours to allow the cells to grow but block cell division. The cultures were then transferred to the dark at 30 °C and the cells were allowed to divide for 24 hours. These partially synchronized cultures were diluted to a cell density of approximately 1x10^7^ cells.ml^-1^ either in ½ SS medium. The cultures were grown at a light intensity of 500 µmol.m^-2^.s^-1^ at 30 °C. The cultures were sampled in the early G1 phase, shortly after one cell cycle was completed.

ChIP-seq experiments were performed with two biological replicates using the photoautotrophically-grown cells described above. ChIP was performed as described previously (Strenkert et al. 2011) with modifications based on (Mozgová et al. 2015). *Chlorella sorokiniana* UTEX1230 cells (9 x 10^8^ cells) were crosslinked at room temperature using 0.37% formaldehyde for 10 minutes, after which crosslinking was quenched by 0.125 M glycine for 10 minutes at room temperature. Cells were washed twice with water and centrifuged at 3000 g for 5 min at 4°C, subsequently resuspended in 200µl of Lysis Buffer (1% SDS, 10 mM EDTA, 50 mM Tris-HCl pH 8, 1x cOmplete™, EDTA-free Protease Inhibitor Cocktail (Roche #04693132001)) and frozen with liquid nitrogen. Next, cells were sonicated using a Bioruptor Plus with 20 cycles of 30 sec ON/OFF at 4°C to achieve DNA fragments size of approximately 200 bp and centrifuged for 5 min at 4500 g at 4°C to remove cell debris. The supernatant was diluted ten times with ChIP dilution buffer (1.1% Triton X-100, 1.2 mM EDTA, 16.7 mM Tris-HCl, pH 8, 167 mM NaCl, cOmplete™, EDTA-free Protease Inhibitor Cocktail (Roche #04693132001)), 200µl aliquot was used for each IP incubated overnight at 4°C with 1.5µg of antibody. Antibodies specific for the following epitopes were used: H3 (Millipore/Merck, 07-690, lot# 3683182), H3K4me3 (Millipore/Merck, 07-473, lot# 3660317), H3K9me2 (Abcam, ab1220, lot #GR3247768-1), H3K27me3 (Diagenode, C15410069, lot # A1818P). The next day, the samples were mixed with 20 µl Dynabeads Protein A (Thermo) and incubated for another 2 hours at 4°C. Next, beads were successively washed three times (5 min/wash) with Low Salt Wash Buffer (150 mM NaCl, 0.1% SDS, 1% Triton X 100, 2 mM EDTA, 20 mM Tris-HCl pH 8), High Salt Buffer (500 mM NaCl, 0.1% SDS, 1% Triton X 100, 2 mM EDTA, 20 mM Tris-HCl pH 8), LiCl Wash Buffer (0.25 M LiCl, 1% NP40, 1% deoxycholate, 1 mM EDTA, 10 mM Tris-HCl pH 8) and finally with TE Buffer (10 mM Tris-HCl pH 8, 1 mM EDTA). DNA was decrosslinked, purified, and eluted using IPureKit (Diagenode). ChIP was performed in three biological replicates, of which two were subject to Illumina sequencing. Sequencing libraries were made with 5 ng DNA using NEBNext® Ultra™ II DNA Library Prep Kit for Illumina (NEB #E7103S/L) following the manufacturer’s instructions. Paired-end 150bp Illumina Sequencing was performed using the NovaSeq 6000 System, with an average output of 20 Mio reads/sample.

## ChIP-seq data analysis

After a survey of existing ChIP-seq data from multiple members of the Viridiplantae, we chose a series of H3K4me3, H3K9me1/2 and H3K27me3 datasets from five Viridiplantae species – the two bryophytes *Physcomitrium patens* and *Anthoceros agrestis* and the three flowering plants *Spirodela polyrhiza*, *Oryza sativa* and *Arabidopsis thaliana* **(Fig. 1A; Supplemental Table S1)**. These were all generated from vegetative tissue of the dominant life cycle phase of each species i.e., haploid gametophytic tissue for the bryophyte species and sporophytic leaf tissue for the flowering plants, and whole plants in case of the duckweed *Spirodela*. For consistency in cross-comparing between different species, we chose not to include *Marchantia polymorpha* given that it was generated with CUT&RUN rather than traditional ChIP-seq (Montgomery et al. 2020). Both the *Chlorella* and public ChIP-Seq data were processed using the Nextflow nf-core/chipseq v2.0.0 pipeline (https://nf-co.re/chipseq/2.0.0/) (Ewels et al. 2020). The ChIP-seq data for each species was mapped to the latest genome assembly and annotation available at the time of our analysis, which are included in an online data repository (https://doi.org/10.17617/3.PJSEUL), together with a MultiQC reports with quality control metrics such as trimming, mapping, coverage and complexity metrics, as well as the version of each tool used in the pipeline. Because the *Physcomitrium* H3K9me2 ChIP-seq data was generated using the legacy SOLiD sequencing format (Widiez et al. 2014), we developed a custom pipeline to process these reads and re-mapped it to the latest *Physcomitrium* patterns genome assembly available at the time of our analysis (https://genomevolution.org/coge/GenomeInfo.pl?gid=33928; see online data repository) using BLAT-like Fast Accurate Search Tool (BFAST v0.7.0a) (https://github.com/nh13/BFAST). First, the reference genome was prepared in color space using the command *bfast fasta2brg* with a *-A 1* flag, then indexed with the command *bfast index* using the *-A 1 -m 1111111111111111111111 -w 14* flag. The color space reads were matched with *bfast match* using a *-A 1 -z -r* flag, then locally aligned with *bfast localalign* using the flag *-A 1 -n 8 -U*. The resulting alignment was then filtered using *bfast postprocess* with the flag *-A 1 -a 2 -O* 1. The downstream processing of the resulting aligned sam file followed that as in the nf-core/chipseq v2.0.0 pipeline. For data visualisation and plotting, normalised log_2_ bigwig coverage files of each histone mark relative to H3 or input were generated using deepTools version 3.5.1 bamCompare with a bin size of 10 bp (Ramírez et al. 2014). Biological replicates were merged where available (**Supplemental Table S1**). A cross-correlation matrix of Spearman’s correlation coefficient of the *Chlorella* samples was generated using deepTools v3.5.1 *multiBamSummary* (Ramírez et al. 2014). Bigwig coverage files were visualized along each genome assembly using IGV v2.16.2 (Robinson et al. 2011).

Functional analysis of H3K27me3-marked genes in *Chlorella* was based on the gene annotations provided by Phycocosm (Chloso_1 assembly). Five different classifications were included: Gene ontology (GO) – biological process (GO_BP), molecular function (GO_MF), and cellular component (GO_CC), Kyoto encyclopedia of genes and genomes (KEGG) – pathway (KEGG_pathway) and pathway class (KEGG_pathway_class) and Eukaryotic orthologous groups (KOG_definition). Hypergeometric tests were performed using the enricher function in the R package clusterProfiler (Yu et al. 2012), with default parameters, considering the five classifications separately. A functional term was considered enriched among the Chlorella H3K27me3-marked genes if its false-discovery rate (FDR) was lower than 5%.

## RNA-seq analysis

The analysis of RNA-seq data from across *Arabidopsis* development was performed as described previously (Borg et al. 2020). In brief, raw FASTQ files for each dataset were downloaded from the Gene Expression Omnibus (GEO) database (**Supplemental Table S1**). Adapters were trimmed using TrimGalore v0.4.1 (https://github.com/FelixKrueger/TrimGalore) and the resulting reads aligned to the *Arabidopsis* genome (TAIR10) using the STAR aligner v2.5.2a (Dobin et al. 2013). Transcripts per million (TPM) values were generated using Kallisto v0.43.1 (Bray et al. 2016) with an index built on TAIR10 cDNA sequences (see online data repository). The *agdp1* and *suvh456* RNA-seq datasets (Zhang et al. 2018) were processed similarly except that read count quantification per gene was summarised using Salmon v1.10.1 in alignment-based mode (Patro et al. 2017). The resulting count table was used for differential gene expression analysis using DESeq2 v1.40.2 (Love et al. 2014). Differentially expressed genes (DEGs) were defined as having a log_2_ fold-change >1 or < -1 and an adjusted *p*-value < 0.05 compared to the wild-type control. Overlap of enrichment of DEGs with *Arabidopsis* TRGs (ranks 8-11) was determined with the R package GeneOverlap v1.36.0 function newGOM (https://github.com/shenlab-sinai/GeneOverlap). Expression specificity *tau* scores of each *Arabidopsis* gene were calculated in R across the RNA-seq datasets shown in **Supplemental Figure S4** as described previously (Lüleci and Yılmaz 2022).

## DNA methylation analysis

Raw FASTQ files for methylomes previously generated from *Physcomitrium patens* (Yaari et al. 2019), *Spirodela polyrhiza* (Harkess et al. 2024), *Oryza sativa* (Xu et al. 2020) and *Arabidopsis thaliana* (Gallego-Bartolomé et al. 2019) were first quality and adapter trimmed with using TrimGalore v0.6.10 with default settings. Bisulfite-converted reads were then aligned against each respective genome in directional mode using Bismark v0.23.0 (*bismark -q –score-min L,0,–0.4*) (Krueger and Andrews 2011). Deduplicated and uniquely mapped alignments were then used to call weighted methylation rates at every converted cytosine using Methylpy v1.2.9 (Schultz et al. 2015). Cytosines with a coverage ≥ 4 were used to calculate the average methylation rate per genomic feature and calculated using the BEDTools v2.30.0 *map* function. For methylation analysis genes or TEs with ≥ 4 mapped cytosines were retained.

## Chromatin state of evolutionarily aged genes

Genomic phylostratigraphy was performed to age protein-coding genes in each species using GenERA with the flag *-u “–query-cover 50”* to increase the threshold of protein homology to at least 50% (Barrera-Redondo et al. 2023). H3K9me1/2- and H3K27me1/3-marked genes were identified using the R packages *ChIPpeakAnno* and *GenomicRanges* (Zhu et al. 2010; Lawrence et al. 2013) and defined as having a minimum overlap of 100bp of coding sequence with the respective ChIP-seq peaks. The log_2_ ratio of observed to expected values in **Figure 2** represents the proportion of genes in each phylogenetic rank marked with a given histone mark (observed) divided by the proportion of genes in each phylogenetic rank found across the whole genome (expected). Circos plots showing the distribution of the phylogenetically-ranked genes in each species were generated using the R package circlize (Gu et al. 2014). Meta profiles of TE enrichment were generated using the EnrichedHeatmap function *normalizeToMatrix* and plotted using a custom script in R. Taxonomically-restricted genes (TRGs) were defined as those that arose within the family and in later ranks for each species (e.g,. those in the Brassicaceae rank onwards in the case of *Arabidopsis*). GO enrichment computation of H3K9me2- and H3K27me3-marked genes in each phylogenetic rank in *Arabidopsis* was performed using Panther 19.0 (DOI: 10.5281/zenodo.12173881).

## Enrichment analysis of H3K27me3-marked *Arabidopsis* genes associated with development

*Arabidopsis* genes were designated H3K27me3-marked as described in the previous section. Developmental genes were defined as annotated with the GO term development (GO:0032502), whereas non-developmental genes were those not associated with this GO term but instead associated with any of the other GO terms from at the same level in the GO tree (GO:0044848 biological phase, GO:0044419 biological process involved in interspecies interaction between organisms, GO:0051703 biological process involved in intraspecies interaction between organisms, GO:0065007 biological regulation, GO:0009987 cellular process, GO:0098754 detoxification, GO:0042592 homeostatic process, GO:0002376 immune system process, GO:0051179 localization, GO:0040011 locomotion, GO:0043473 pigmentation, GO:0050896 response to stimulus, GO:0048511 rhythmic process, and GO:0050789 regulation of biological process). Functional annotation of genes was based on the Bioconductor annotation package *org.At.tair.db 3.17.0* (IEA – inferred from electronic annotation, NAS – non-traceable author statement, and ND – no biological data available excluded – Supplementary Table S4). Fisher’s enrichment test was performed in R to assess whether developmental genes were significantly enriched among H3K27me3-marked genes as opposed to non-developmental genes.

## TE annotation and divergence landscapes

*De novo* annotation of TEs and TE library preparation, including the consensus sequences of all identified TE families, was performed for all six Viridiplantae species using EDTA v2.0.0 (Ou et al. 2019). Each TE library was used to mask TEs across the respective genomes using RepeatMasker vopen-4.0.9 (Smit, AFA, Hubley, R & Green, P. *RepeatMasker Open-4.0*. 2013-2015 http://www.repeatmasker.org). To identify repeats and TEs marked with H3K9me1/2, H3K27me3 or both, ChIP-seq peaks were intersected with each repeat or TE using the R packages *ChIPpeakAnno* and *GenomicRanges* (Zhu et al. 2010; Lawrence et al. 2013). TE landscapes of DNA transposons, LTR retrotransposons and MITEs were generated from the RepeatMasker output file (.out file) using the percent diversity parameter, which represents the percentage of substitutions of a given TE relative to the consensus sequence of its assigned family. Given that EDTA is unable to successfully annotate LINE elements (Ou et al. 2019), this class of TEs were not considered in our study and thus warrant further investigation in the future.

## Analysis of conserved PRC2-regulated gene families

Orthology inference analysis was first performed to identify orthologous gene families (or orthogroups) across the proteome of each species using OrthoFinder v2.5.2 (Emms and Kelly 2019). The relative proportion of genes marked with H3K27me3 within each orthogroup was then computed for each species to identify conserved PRC2 target genes. The shared land plant PRC2 network of genes was defined as having at least one gene within an orthogroup marked with H3K27me3 in each of the five land plant species. The deeply-conserved PRC2-regulated network of genes was similarly defined but with the inclusion of orthogroups that were also conserved in *Chlorella*, whereas the flowering-plant specific PRC2- regulated network was restricted to orthogroups that has at least one gene marked with H3K27me3 exclusively in *Spirodela*, *Oryza* or *Arabidopsis*. Cochran’s Q test and binomial tests were performed in R. The hypothesised probability of success (p) of the binomial tests was based on all observed proportions of H3K27me3-marked orthogroups to reflect the differing variability of H3K27me3 marking in each species. Functional annotation of the orthogroups was performed with *Arabidopsis* gene identifiers using the online database GenFAM (https://www.mandadilab.com/genfam/) (Bedre and Mandadi 2019). Further manual curation was performed to assign the orthogroups to one of the nine functional classes defined in **Figure 5C**. The annotation of transcription-associated proteins (TAPs) was performed with the proteome of each species using TAPscan v4 (https://tapscan.plantcode.cup.uni-freiburg.de) (Petroll et al. 2024).

## Motif enrichment analysis

Motif enrichment was performed using motifs in the DAP-seq data set (O’Malley et al. 2016) with the *AME* function in the MEME suite version 5.5.4 (Bailey et al. 2009). The H3K27me3 peaks overlapping genes that form part of the shared land plant PRC2-regulated network in each species were provided as input, with a shuffled set of peaks used as the background model. To rule out false-positive motifs, we also performed motif enrichment analysis on conserved non-H3K27me3 target genes using all genes in each genome as the background model. Commonly enriched motifs were defined as having an adjusted *p*-value less than 0.05 in each land plant species that were filtered for the false-positive motifs. Heatmaps of DAP-seq peaks, motif occurrences and H3K27me3 enrichment were generated using the R package EnrichedHeatmap (Gu et al. 2018).

## Data availability

*Chlorella* ChIP-seq data generated in this study have been deposited in the European Nucleotide Archive (ENA) under accession code PRJEB79967. A repository containing all the genome assemblies, annotation and processed data generated in this study can be accessed at https://doi.org/10.17617/3.PJSEUL, which includes re-analysis of the public datasets listed in **Supplemental Table S1**. All scripts are available upon request.

## Contributions

MB conceived and designed the study. MB performed the bioinformatic analyses with support from RP, RKP and AL. Green algal histone H3 sequences were collated and analysed by MB and IM. Synchronised *Chlorella* cells in G1 were prepared by KB. *Chlorella* ChIP-seq data was generated by RK under supervision of IM. QR and IM performed the functional enrichment analysis of H3K27me3 target genes. RP generated the TE landscapes for Figure 4. RKP processed public DNA methylation datasets and the *adgp1* and *suvh456* RNA-seq data for Supplemental Figures 2 and 3. AL surveyed public ChIP-seq datasets and developed scripts for downstream analysis under supervision of MB. MB interpreted the data, assembled the figures and wrote the manuscript.

## Acknowledgements

The authors thank Josue Barrera-Redondo and Sodai Lotharukpong for advice on running GenERA. MB, RP and AL were supported by the Max-Planck-Gesellschaft. RK was supported by an ERA Fellowship from the European Research Executive Agency (101090308 – PAMFGAL). IM was supported by an ERC-CZ grant from the Czech Academy of Sciences (ERC200961901). Computational resources to MB group were provided by the Max Planck Institute for Biology and for IM group were provided by the e-INFRA CZ project (ID:90254), supported by the Ministry of Education, Youth and Sports of the Czech Republic.

---

**Supplemental Figure S1.**
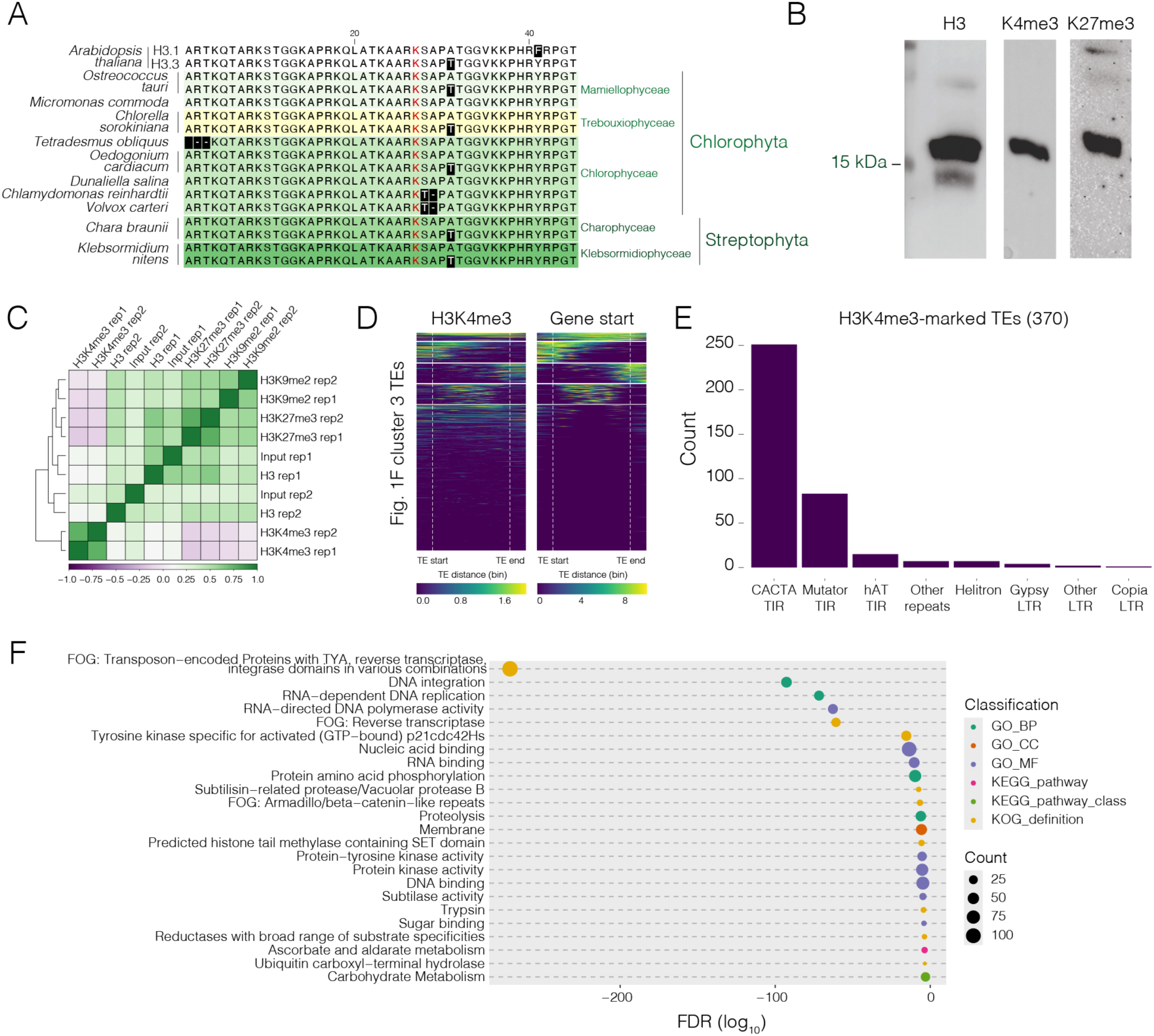
(A) ClustalW alignment of the N-terminal tail of histone H3 variants among various clades of chlorophyte and streptophyte green algae. *Arabidopsis* histone H3.1 and H3.3 are included as an outgroup. Lysine 27 (K27) is highlighted in red. (B) Western blot analysis of H3, H3K4me3 and H3K27me3 in *Chlorella sorokiniana*. (C) Spearman cross-correlation matrix of the *Chlorella* ChIP-seq data sets generated in this study. (D) A metaplot of H3K4me3 enrichment and gene start signals over cluster 3 TEs from Figure 1F. “Gene start”: Regions +/- 50 bp around a TSS are assigned a score of 10 and are penalized by -1 for each 50bp of increasing distance from a TSS. Thus, genomic regions outside a -500 to +500 bp interval around a TSS are assigned with a 0 score. (E) Classification of the 370 TE models overlapping an H3K4me3 peak. (F) Enriched functional terms among the H3K27me3-marked genes in *Chlorella* (FDR < 0.05, hypergeometric test). Terms are plotted with their FDR (in log10) with dots coloured according to their classification and sized according to their count (number of genes among the H3K27me3-marked genes). GO_BP: Gene ontology, biological process; GO_MF: Gene ontology, molecular function; GO_CC: Gene ontology, cellular component; KEGG_pathway: Kyoto encyclopedia of genes and genomes, pathway; KEGG_pathway_class: Kyoto encyclopedia of genes and genomes, pathway class; KOG_definition: Eukaryotic orthologous groups, definition.

**Supplemental Figure S2.**
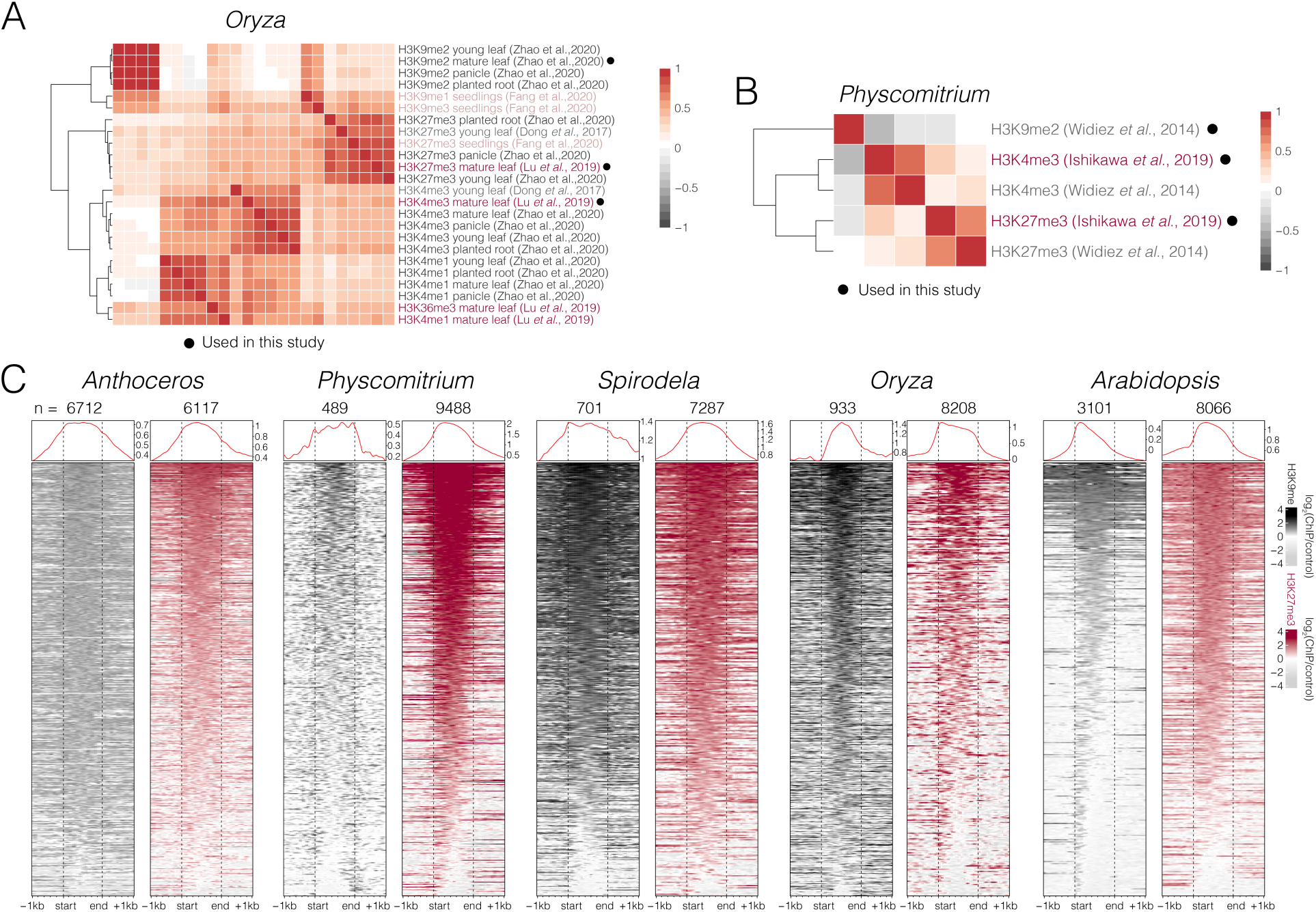
(A-B) Pearson cross-correlation matrix of ChIP-seq data from *Oryza sativa* (A) and *Physcomitrium patens* (B) from independent studies. The black dots indicate the final data sets used in this study. (C) Levels of H3K9me2 (or H3K9me1 in *Anthoceros*) and H3K27me3 marks centred over genes overlapping ChIP-seq peaks of each respective form of histone methylation in each land plant species (see Supplemental Table S3). Plotted is the log_2_ ChIP-seq enrichment relative H3 or input in the case of H3K9me2 in Spirodela and Oryza. Numbers of genes overlapping either form of histone methylation are indicated above each heatmap.

**Supplemental Figure S3.**
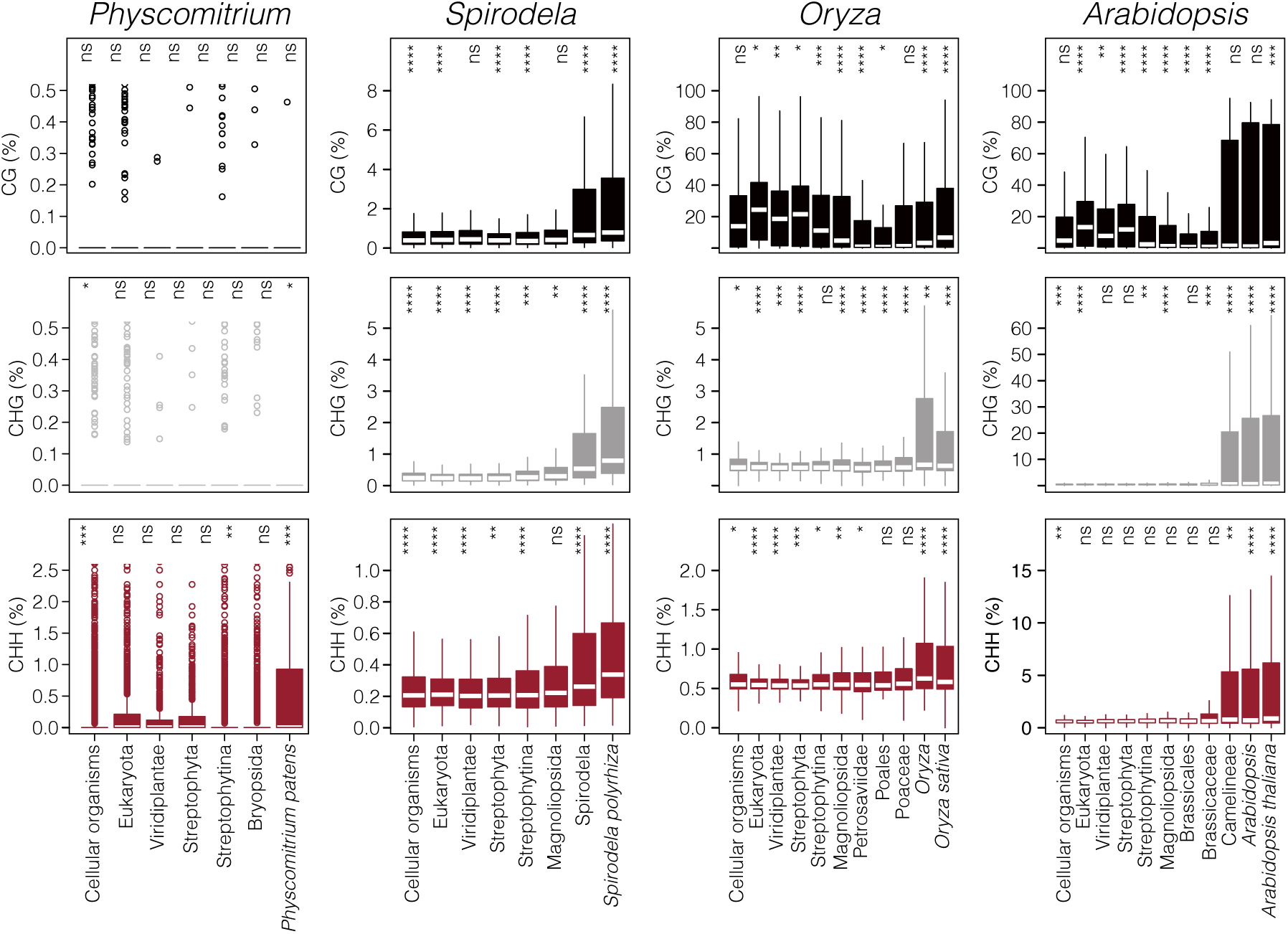
Boxplots depicting the weighted CG, CHG, and CHH methylation rates (top to bottom) across phylogenetically-ranked genes in *Physcomitrium patens*, *Spirodela polyrhiza*, *Oryza sativa* and *Arabidopsis thaliana* (left to right). Thick horizontal bars mark the medians, box edges correspond to the 25th and 75th percentiles, and whiskers extend to 1.5× the interquartile range (IQR). Significance *p*-values were computed using a Wilcoxon test against the mean methylation rate of all genes per species.

**Supplemental Figure S4.**
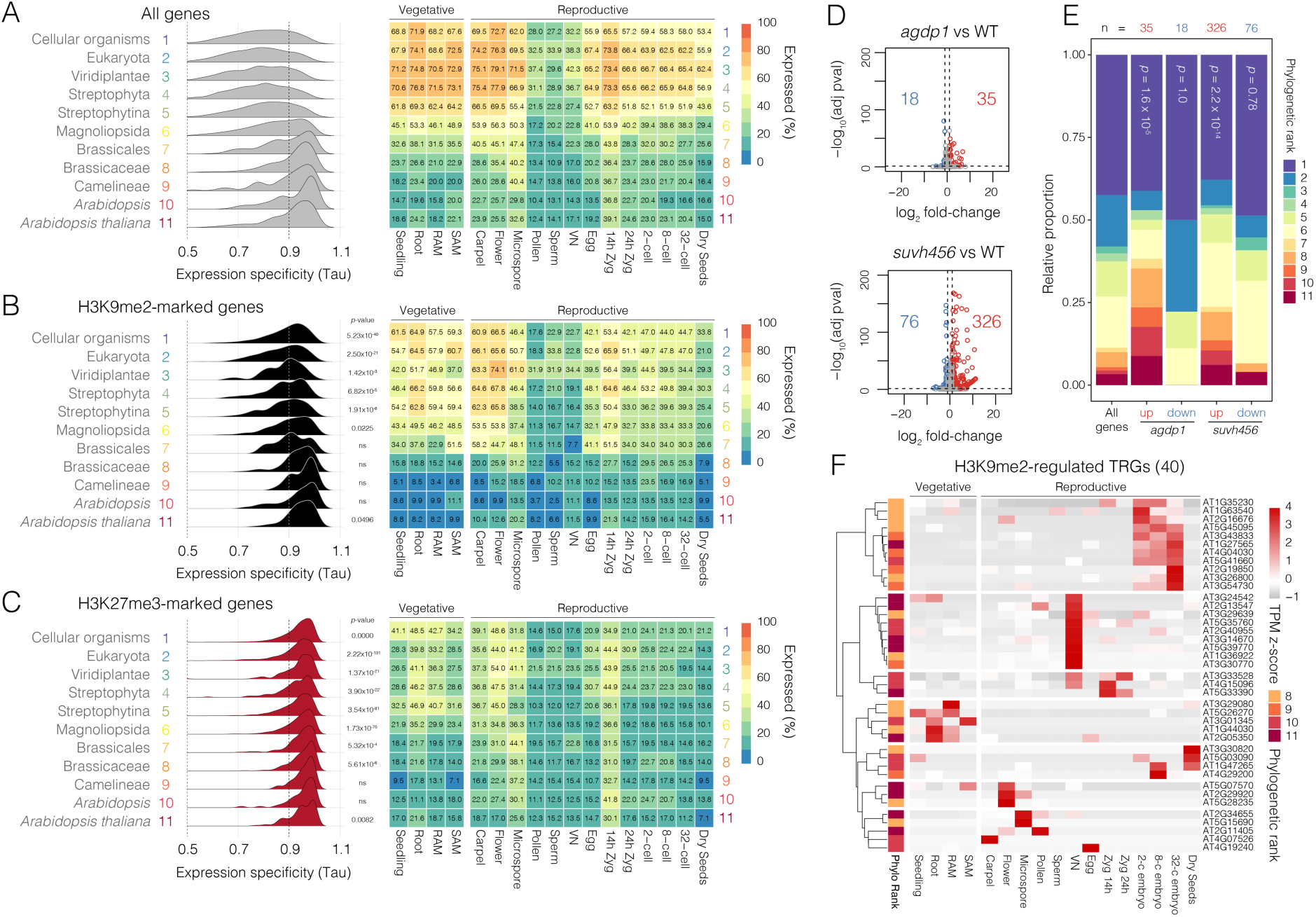
(A-C) Ridgeline plots (left) showing the distribution of tau values for all phylogenetically-ranked genes in Arabidopsis (A) and those marked with H3K9me2 (B) and H3K27me3 (C). High *tau* > 0.9 (dashed line) indicates genes with highly restricted expression in the different tissues shown in the heatmaps to the right. The heatmaps summarise the percentage of genes in each phylogenetic rank that are expressed in a given tissue or cell type (TPM > 5). Significant p-values indicated on the ridgeline plots of panels B and C are the result of a Mann-Whitney U test comparing the distribution of tau values in with those of all genes in the genome for each phylogenetic rank (panel C). ns = no significant difference. (D) Volcano plot of differentially expressed genes in *agdp1* or *suvh456* mutants compared to wild type (WT) *(Zhang et al. 2018)*. Upregulated genes are shown in red while downregulated genes are shown in blue. DEGs were defined as having a log_2_ fold-change > 1 and FDR-adjusted p-value < 0.05. (E) Relative proportion of phylogenetically-ranks among all Arabidopsis genes and among the DEGs in panel D. The p-value indicates the statistical enrichment of taxonomically-restricted genes (TRGs; ranks 8-11) among each group of DEGs as determined using pairwise two-sided Fisher’s exact tests. (F) Heatmap illustrating the developmentally-regulated expression of H3K9me2-regulated TRGs (i.e., H3K9me2- marked TRGs that are also upregulated in *suvh456* mutants compared to WT). Expression represents the z-score of normalised RNA-seq TPM values. The left panel indicates the phylogenetic rank assigned to each TRG.

**Supplemental Figure S5.**
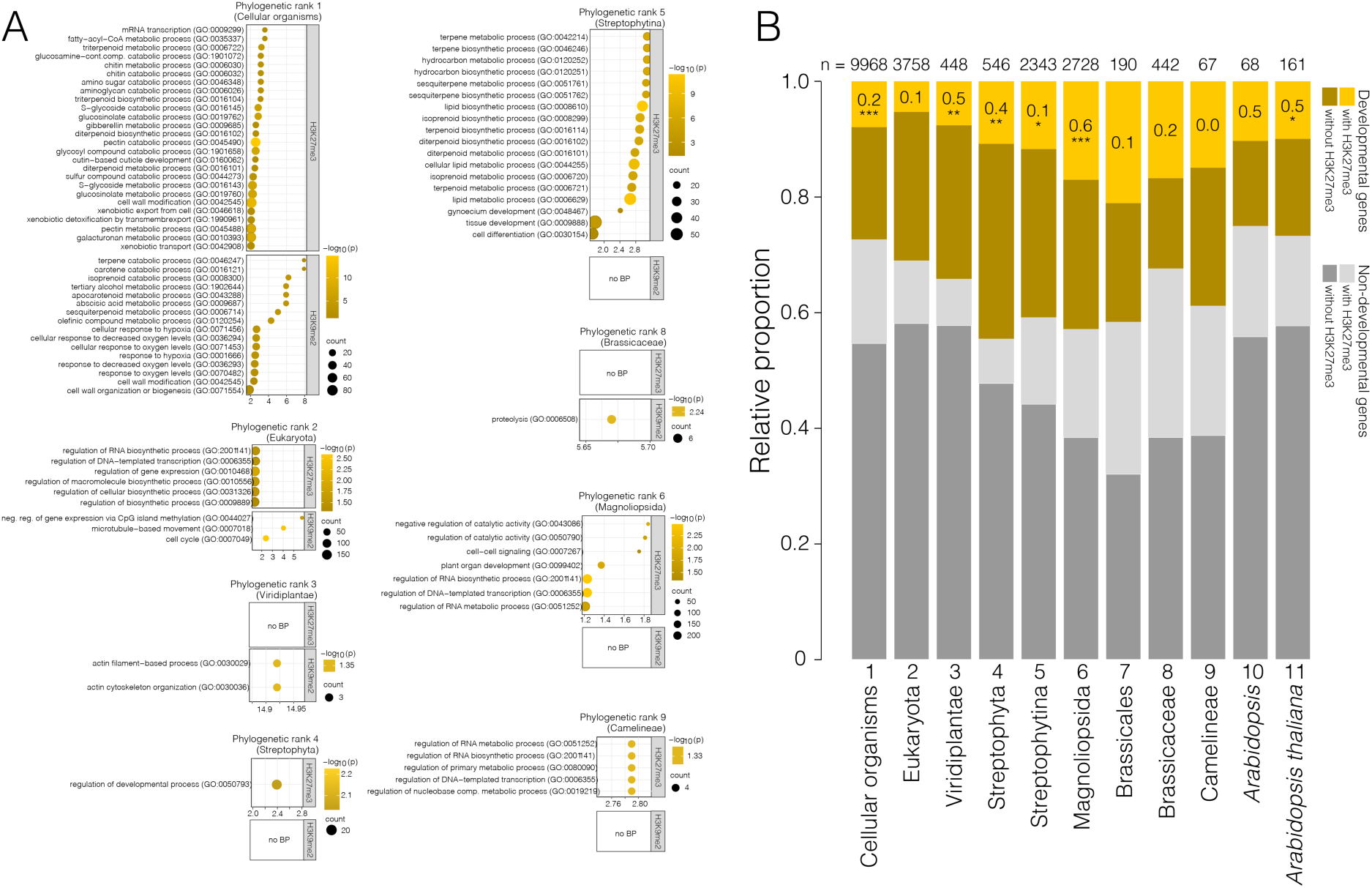
(A) Gene ontology (GO) enrichment analysis of H3K27me3-marked and H3K9me2-marked genes in each of the 11 phylogenetic ranks in *Arabidopsis*. All genes assigned to the corresponding phylogenetic rank were used as the reference gene set for the computation of *p*-values. Enriched GO terms belonging to the Biological Processes (BP) category are shown, with only those enriched more than 2-fold displayed for H3K27me3 targets in Phylogenetic rank 1 (See Supplemental Table S4 for more terms). Phylogenetic ranks 7, 10 and 11 with no significant enrichment of BP categories were not plotted. (B) Partitioning of phylogenetically-ranked *Arabidopsis* genes by H3K27me3 marking and association with the GO term “Development” (GO:0032502). Asterisks indicate significance based on a Fisher’s test of the enrichment of “developmental” genes among H3K27me3-marked genes (blue) as opposed to the “non-developmental” genes (red). * (*p*-value < 0.05), ** (*p*-value < 0.01) or *** (*p*-value < 0.001). Log_2_ fold-enrichment is indicated in the top sector, while the number of genes analysed in each phylogenetic rank are indicated above the stacked bars (n).

**Supplemental Figure S6.**
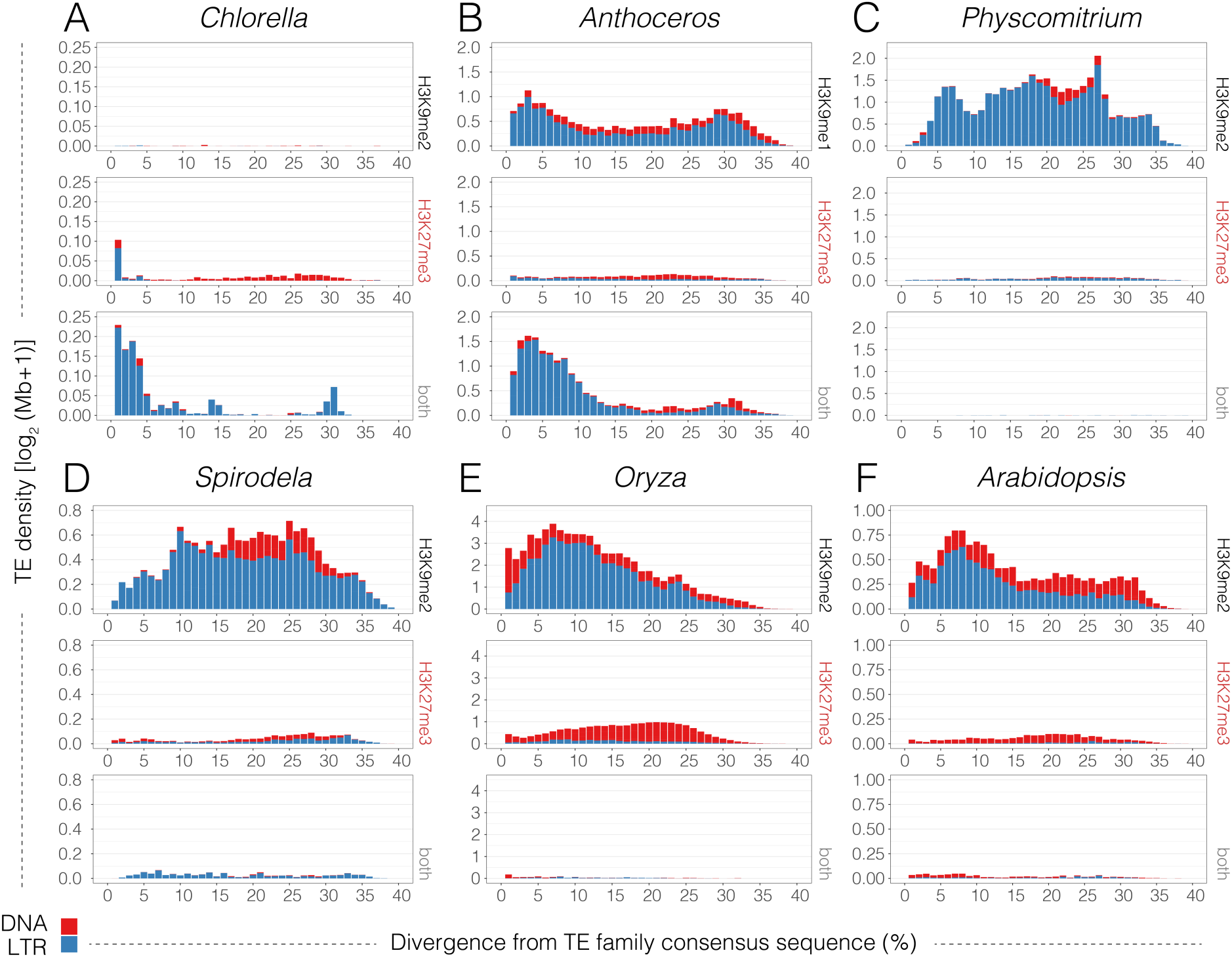
Histograms summarising the relative age and distribution of DNA (red) and LTR (blue) transposons marked with H3K9me only (top row), H3K27me3 only (middle row) or both (bottom row) in *Chlorella* (A), *Anthoceros* (B), *Physcomitrium* (C), *Spirodela* (D), *Oryza* (E) and *Arabidopsis* (F). Increasing sequence divergence of TEs from the consensus sequence of their assigned family (x-axis) is indicative of an older age and reduced potential for transposition.

**Supplemental Figure S7.**
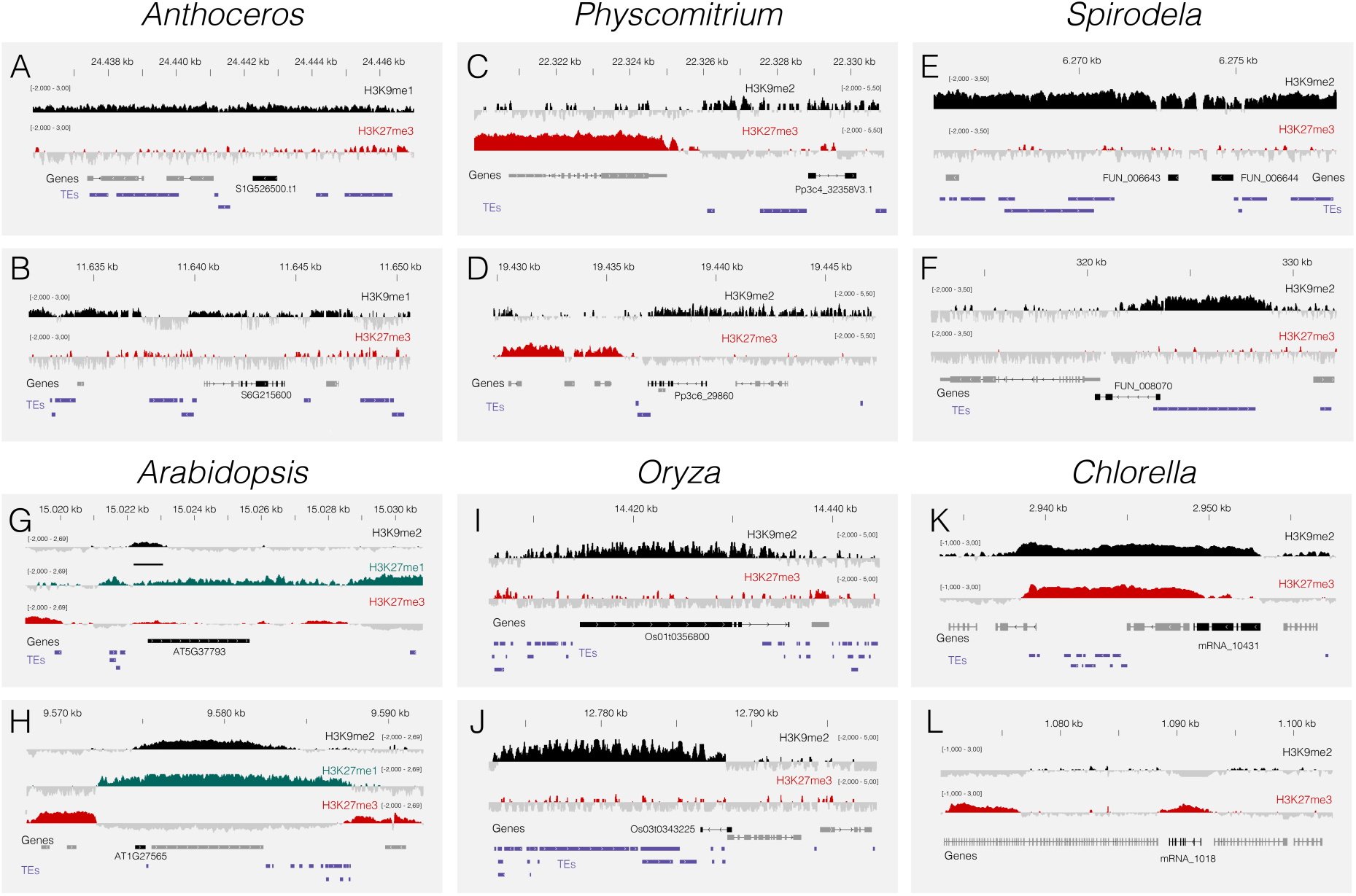
ChIP-Seq tracks showing H3K9me2 (black; H3K9me1 in *Anthoceros*) and H3K27me3 (red) levels over TRGs in *Anthoceros* (A-B), *Physcomitrium* (C-D), *Spirodela* (E-F), *Arabidopsis* (G-H), *Oryza* (I-J) and *Chlorella* (K-L). Coloured and grey shading indicate enriched or depleted signals, respectively. Coverage is represented as the ChIP-seq log_2_ ratio relative to H3 (range is indicated in brackets). Gene models are shown below in grey while the TRG-of-interest is highlighted in black alongside its gene ID, with TEs shown in purple.

**Supplemental Figure S8.**
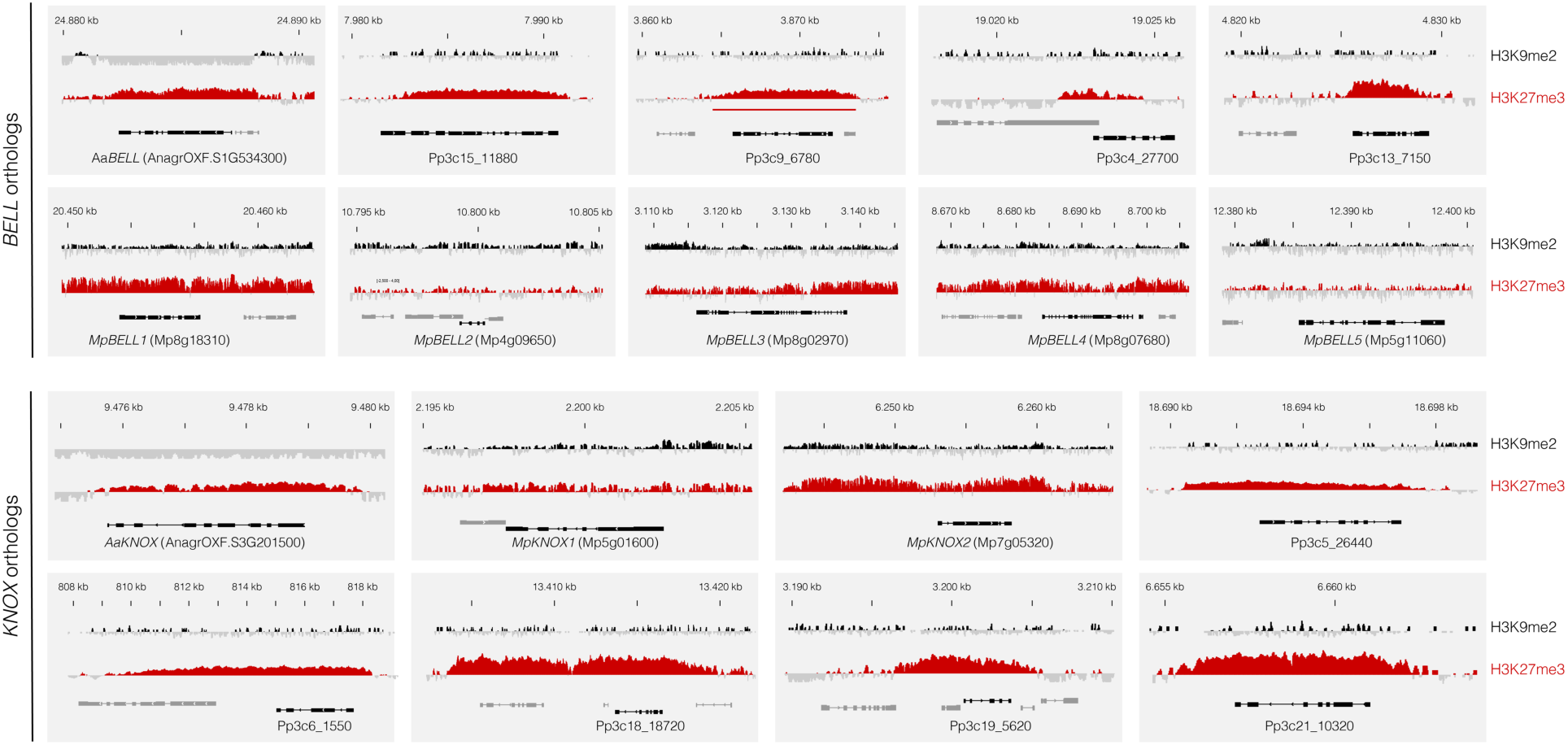
ChIP-Seq tracks showing H3K9me2 (black; H3K9me1 in *Anthoceros*) and H3K27me3 (red) levels over BELL and KNOX orthologs in *Anthoceros* (Aa), *Marchantia* (Mp) and *Physcomitrium* (Pp). Coloured and grey shading indicate enriched or depleted signals, respectively. Coverage is represented as the ChIP-seq log_2_ ratio relative to H3.

## Supplemental Tables

**Supplemental Table S1**. Publicly available genomic datasets used in this study.

**Supplemental Table S2**. Statistical summary in support of main Figure 2.

**Supplemental Table S3**. Phylogenetic rank and chromatin state of the genes from each species.

**Supplemental Table S4**. Enriched GO terms among the H3K9me2- and H3K27me3-marked genes of differing evolutionary age in *Arabidopsis*.

**Supplemental Table S5**. Orthogroups identified in this study.

**Supplemental Table S6**. H3K27me3-marked orthogroups.

**Supplemental Table S7**. Shared land plant PRC2-regulated network.

**Supplemental Table S8**. Deeply-conserved PRC2-regulated network.

**Supplemental Table S9**. Transcription-associated proteins regulated by PRC2 across the green lineage.

**Supplemental Table S10.** DAP-seq motifs enriched within the H3K37me3 domains marking genes in the shared land plant PRC2-regulated network.

